# Iron is a ligand of SecA-like metal-binding domains *in vivo*

**DOI:** 10.1101/613315

**Authors:** Tamar Cranford-Smith, Mohammed Jamshad, Mark Jeeves, Rachael A. Chandler, Jack Yule, Ashley Robinson, Farhana Alam, Karl A. Dunne, Edwin H. Aponte Angarita, Mashael Alanazi, Cailean Carter, Ian R. Henderson, Janet E. Lovett, Peter Winn, Timothy Knowles, Damon Huber

## Abstract

The ATPase SecA is an essential component of the bacterial Sec machinery, which transports proteins across the cytoplasmic membrane. Most SecA proteins contain a long C-terminal tail (CTT). In *Escherichia coli*, the CTT contains a structurally flexible linker domain and a small metal-binding domain (MBD). The MBD coordinates zinc via a conserved cysteine-containing motif and binds to SecB and ribosomes. In this study, we screened a high-density transposon library for mutants that affect the susceptibility of *E. coli* to sodium azide, which inhibits SecA-mediated translocation. Results from sequencing this library suggested that mutations removing the CTT make *E. coli* less susceptible to sodium azide at subinhibitory concentrations. Copurification experiments suggested that the MBD binds to iron and that azide disrupts iron binding. Azide also disrupted binding of SecA to membranes. Two other *E. coli* proteins that contain SecA-like MBDs, YecA and YchJ, also copurified with iron, and NMR spectroscopy experiments indicated that YecA binds iron via its MBD. Competition experiments and equilibrium binding measurements indicated that the SecA MBD binds preferentially to iron and that a conserved serine is required for this specificity. Finally, structural modelling suggested a plausible model for the octahedral coordination of iron. Taken together, our results suggest that SecA-like MBDs likely bind to iron *in vivo*.

## INTRODUCTION

Translocation of proteins across the cytoplasmic membrane is carried out by the evolutionarily conserved Sec machinery (1,2). The central component of this machinery is a channel in the cytoplasmic membrane, which is composed of the proteins SecY, -E and -G in bacteria (3). The translocation of a subset of proteins through SecYEG requires the activity of the ATPase SecA (4,5). In *Escherichia coli*, SecA contains 901 amino acids. The N-terminal ∼830 amino acids of SecA in *E. coli* make up the catalytic core of the protein, which is essential for viability and is sufficient to reconstitute protein translocation *in vitro* (6-9). The C-terminal ∼70 amino acids (C-terminal tail; CTT) are widely, but not universally, conserved (10). The CTT contains two regions: a structurally flexible linker domain (FLD) and a small metal-binding domain (MBD). The amino acid sequence of the FLD is poorly conserved, and recent research suggests that the FLD autoinhibits the activity of SecA by stabilising a conformation of SecA with a lower affinity for substrate protein (10,11). When present in SecA, the sequence of the MBD is highly conserved (10). The MBD interacts with SecB and ribosomes (10,12), and interaction of the MBD with SecB or ribosomes relieves MBD-mediated autoinhibition (10,11). The MBD binds to a single Zn^2+^ ion (12), which is coordinated by three conserved cysteines and a histidine with a conserved CXCX_8_CH (CXCX_8_CC in some bacteria) motif (**supporting figure S1**) (10,12,13). In addition, the MBD contains several highly conserved amino acids, including Ser-889 and Tyr-893, whose functional significance is unknown (10,13,14).

Sodium azide is a well studied inhibitor of SecA (15-18). Azide causes a nearly complete block in SecA-mediated protein translocation within minutes of addition to growing cells (15), and mutations that confer increased resistance to azide map to the *secA* gene (15-17). Azide is thought to inhibit nucleotide exchange by SecA (19,20). Indeed, most of the mutations that confer azide resistance modify amino acids in one of the two nucleotide binding domains (17), and many of these substitutions increase the rate of nucleotide exchange in *in vitro* assays (20). However, the concentration of azide required to partially inhibit the rate of ATP turnover by SecA *in vitro* (10-20 mM) is much higher than that required block translocation *in vivo* (1-2 mM) (15,21), suggesting that azide-mediated inhibition of SecA *in vivo* is more complex.

In order to gain insight into the effect of azide on protein translocation *in vivo*, we determined the effect of azide on the relative growth rates of >1 million independent transposon mutants using <u>tra</u>nsposon-<u>d</u>irected insertion-site <u>s</u>equencing (TraDIS) (22,23). The results of this screen suggested that azide caused CTT-mediated autoinhibition of SecA, potentially by disrupting the MBD. Treating cells with azide reduced the amount of iron (but not zinc) that co-purified with the MBD to background levels, suggesting that a subpopulation of SecA proteins bind to iron and that azide triggers autoinhibition by disrupting the MBDs of this subpopulation. Full-length SecA produced at more physiological levels copurified predominantly with iron. Furthermore, two *E. coli* proteins containing SecA-like MBDs, YecA and YchJ, bind to and copurify with iron. Biophysical characterisation of metal binding properties of YecA and SecA indicated that their MBDs bind to iron, and competition experiments and equilibrium binding measurements suggest that the MBD of SecA binds preferentially to iron. Finally, our results suggest that an evolutionarily conserved serine residue is important for mediating this specificity. Taken together, our results indicate that iron is a physiological ligand of SecA-like MBDs.

## RESULTS

### Identification of genes that affect the susceptibility of E. coli to sodium azide

To gain insight into the effect of azide on cells, we identified transposon mutations that influence the susceptibility of *E. coli* to azide using TraDIS (22,23). To this end, we determined the concentration of azide that inhibited, but did not prevent, the growth of *E. coli* BW25113. Growth curves in the presence of increasing concentrations of azide indicated that 250 and 500 μM sodium azide were the highest concentrations that did not completely inhibit growth in LB (**supporting figure S2**). We then grew a library of > 1 million independent BW25113 mini-Tn*5* transposon mutants (23) in the presence of subinhibitory concentrations of azide and determined the change in the relative proportion of mutations in each insertion using Illumina sequencing. Growth of the library in the presence of 0.25 mM or 0.5 mM azide resulted in a strong depletion (> log_2_ 1.5-fold) in insertions in approximately 180 of the 4346 genes included in the analysis. Many of the most strongly depleted insertions disrupted genes encoding components of the Sec machinery (e.g. *yajC, secG, secF, secM, yidC*) or cell envelope stress responses (e.g. *cpxR, dsbA*) (**figure 1A; supporting data S1**), consistent with the ability of azide to inhibit Sec-dependent protein translocation and disrupt biogenesis of the cell envelope.

**Figure 1.**
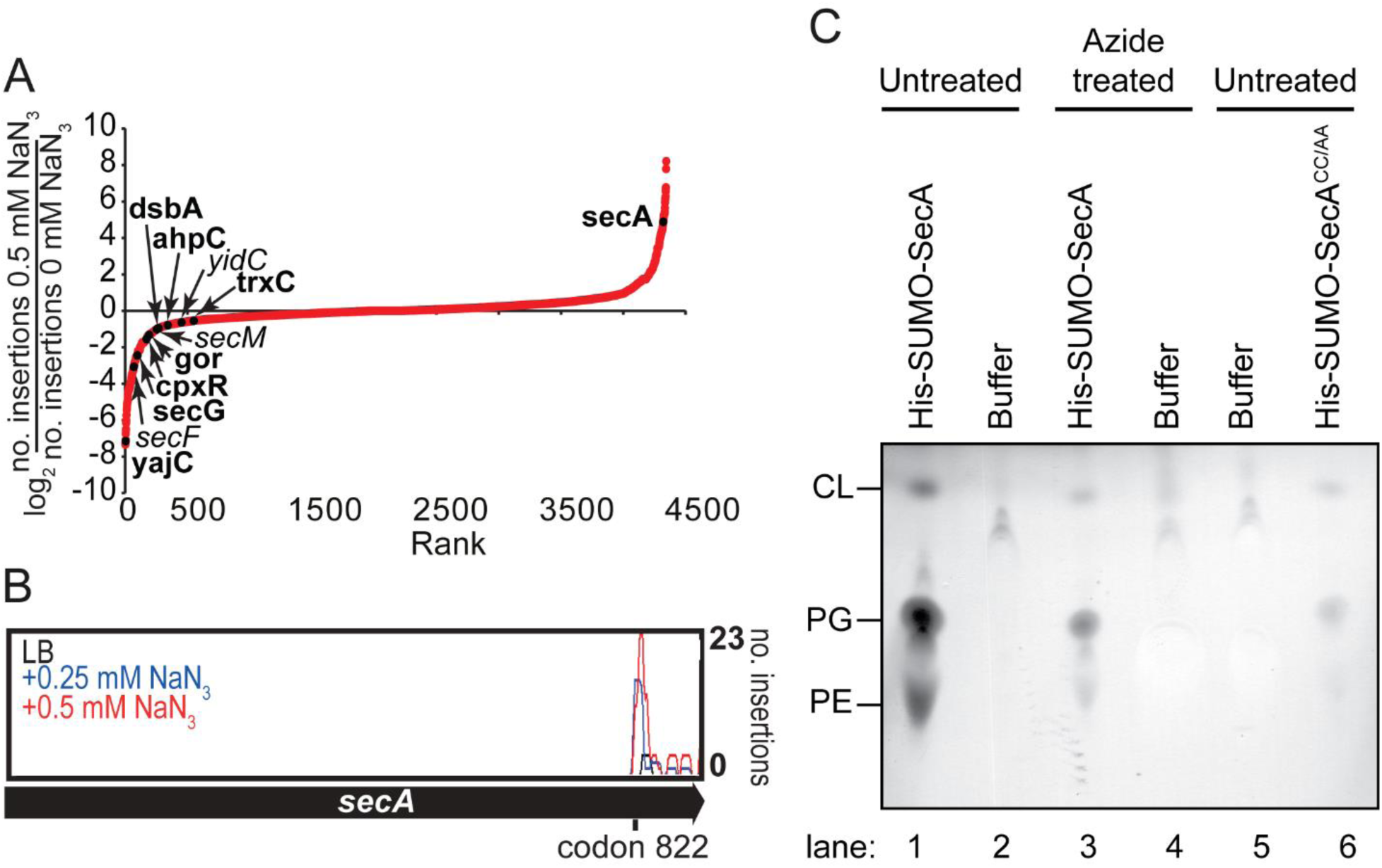
Effect of azide on the C-terminal tail of SecA. (A) Plot of the degree of depletion or enrichment in transposon insertions in all of the non-essential genes in *E. coli* after growth of the TraDIS library to OD_600_ = 0.9 in the presence of 0.5 mM sodium azide. The log_2_ of the fraction of the number of insertions in a gene after growth in the presence of 0.5 mM azide over the number of insertions after growth in LB was plotted according to the degree of enrichment. Data points representing insertions in the *secA, secF, secG, secM, yidC, yajC, cpxR, dsbA, ahpC, gor* and *trxC* genes are indicated. Single gene deletion mutants that were compared to growth of the parent in **table 1** are indicated (bold). (B) Number of mutants containing insertions at the indicated location in the *secA* gene after growth of the TraDIS library in the absence (black) or presence of 0.25 (blue) or 0.5 mM (red) NaN_3_. Most of these insertions truncate the *secA* gene between codons 822 and 829 at the junction between the HSD and the CTT. (C) Cultures of *E. coli* producing His-SUMO-SecA (lanes 1-4) or His-SUMO-SecA^CC/AA^ (lanes 5 & 6) were grown to late exponential phase. In the case of His-SUMO-SecA, half of the culture was treated with 2 mM sodium azide for 10 minutes prior to lysis (lanes 3 & 4), and the other half was left untreated (lanes 1 & 2). His-SUMO-SecA and SUMO-SecA^CC/AA^ were purified from the cell lysates using Ni-affinity purification. Phospholipids from 2 mg of the purified protein were extracted into 100 μl chloroform, and 5μl of the extracted phospholipids (lanes 1, 3 and 6) and the wash buffer (lanes 2, 4 and 5) were resolved using thin-layer chromatography (TLC). The positions of phosphatidylglycerol (PG), phosphatidylethanolamine (PE) and cardiolipin (CL) are indicated.

To verify the results of the TraDIS screen, we investigated the effect of subinhibitory azide on individual mutants. Mutants containing insertions in the *secG, yajC, dsbA, cpxR, gor, ahpC* and *trxC* genes were strongly to moderately depleted during growth in the presence of azide. With the exception of the Δ*gor* mutant, cells containing deletion mutations in these genes grew more slowly than the parent in the presence of azide (**table 1**).

**Table 1.** Relative growth rates of single gene deletion mutants in the absence and presence of sodium azide.

| Strain | Growth rates* |  |  |
| --- | --- | --- | --- |
|  | LB | 0.25 mM NaN<sub>3</sub> | 0.5 mM NaN<sub>3</sub> |
| BW25113 | 1.00 ± 0.00 | 1.00 ± 0.09 | 1.00 ± 0.09 |
| *ΔyajC* | 0.88 ± 0.05 | 0.59 ± 0.08 | 0.54 ± 0.11 |
| *ΔsecG* | 1.02 ± 0.05 | 0.53 ± 0.11 | 0.75 ± 0.1 |
| *ΔdsbA* | 1.01 ± 0.08 | 0.95 ± 0.09 | 0.63 ± 0.15 |
| *ΔcpxR* | 1.13 ± 0.13 | 0.93 ± 0.02 | 0.65 ± 0.15 |
| *Δgor* | 0.96 ± 0.01 | 0.97 ± 0.08 | 1.02 ± 0.04 |
| *ΔahpC* | 0.95 ± 0.00 | 0.80 ± 0.06 | 0.64 ± 0.06 |
| *ΔtrxC* | 0.92 ± 0.00 | 0.89 ± 0.02 | 0.78 ± 0.04 |
| MC4100 | 1.00 ± 0.02 | ND<sup>†</sup> | 1.00 ± 0.03 |
| *secA827::IS1* | 1.03 ± 0.01 | ND<sup>†</sup> | 1.38 ± 0.18 |

\*Parent and mutants were grown in LB or LB containing the indicated concentration of sodium azide.
Growth rates were determined by measuring the optical density at 600 nm every 30 minutes and normalised to the rate of growth of the parent strain under the same conditions.
<sup>†</sup>not determined

The presence of azide also caused a strong enrichment in mutants containing insertions in the *secA* gene at codon 822 (**figure 1B**), consistent with the requirement for the N-terminal portion of SecA for viability. Growth in the presence of azide resulted in a strong enrichment in insertions truncating SecA between amino acids 822 and 829, resulting in a complete truncation of the CTT (**figure 1B**). In addition, a strain containing a mutation that truncates SecA at amino acid ∼827 (*secA827*::IS*3*) (6) grew more rapidly than the parent strain in exponential phase in the presence of sodium azide (**table 1**). Despite the increased growth rate at subinhibitory concentrations of azide, this mutant did not detectably influence the MIC of azide in filter disc assay (data not shown). These results suggested that azide causes autoinhibition of SecA at concentrations that are subinhibitory for growth.

### Effect of azide on binding of SecA to membrane phospholipids

We next investigated the effect of azide on binding of SecA to membranes. Previous studies indicate that SecA binds strongly to membranes (24-28), and a protein fusion between hexa-histidine-tagged SUMO and SecA (His-SUMO-SecA) copurified strongly with membrane phospholipids (**figure 1C**). Expression of this fusion protein can complement the conditional growth defect of a strain producing SecA under control of an arabinose-inducible promoter, indicating that it is functional *in vivo*. Previous studies have suggested that the CTT is involved in binding phospholipids (29). Indeed, His-SUMO-SecA containing alanine substitutions in two of the metal-coordinating cysteines (C885 and C887; His-SUMO-SecA^CC/AA^) did not copurify with phospholipids (**figure 1C**). His-SUMO-SecA purified from cells treated with azide prior to lysis did not copurify with phospholipids, indicating that azide disrupts binding of SecA to the membrane (**figure 1C**). Furthermore, washing purified His-SUMO-SecA disrupted binding to phospholipids *in vitro* (**supporting figure S3**). These results suggested that azide could disrupt the structure of the MBD.

### Effect of azide on the MBD of SecA

To investigate the ability of azide to disrupt the structure of the MBD, we determined the metal content of a fusion protein between SUMO and the CTT (SUMO-CTT). ICP-MS indicated that SUMO-CTT copurified with significant amounts of zinc, iron and copper. Treating cells with azide prior to lysis did not affect the amount of zinc or copper that copurified with SUMO-CTT. However, there was a strong reduction in the amount of iron that copurified with SUMO-CTT from azide treated cells (**figure 2A**), raising the possibility that iron is the physiological ligand of the MBD. In addition, SUMO-CTT purified from azide-treated cells was more prone to aggregation than SUMO-CTT purified from untreated cells (**supporting figure S4**), consistent with the notion that azide treatment disrupts the structure of the MBD. Finally, the presence of the metal-chelator EDTA in the growth medium increased the sensitivity of *E. coli* to azide, while the addition of excess FeSO_4_, but not ZnSO_4_, to the growth media decreased sensitivity to azide (**supporting table S1**), supporting the idea that azide disrupts the iron-bound form of SecA.

**Figure 2.**
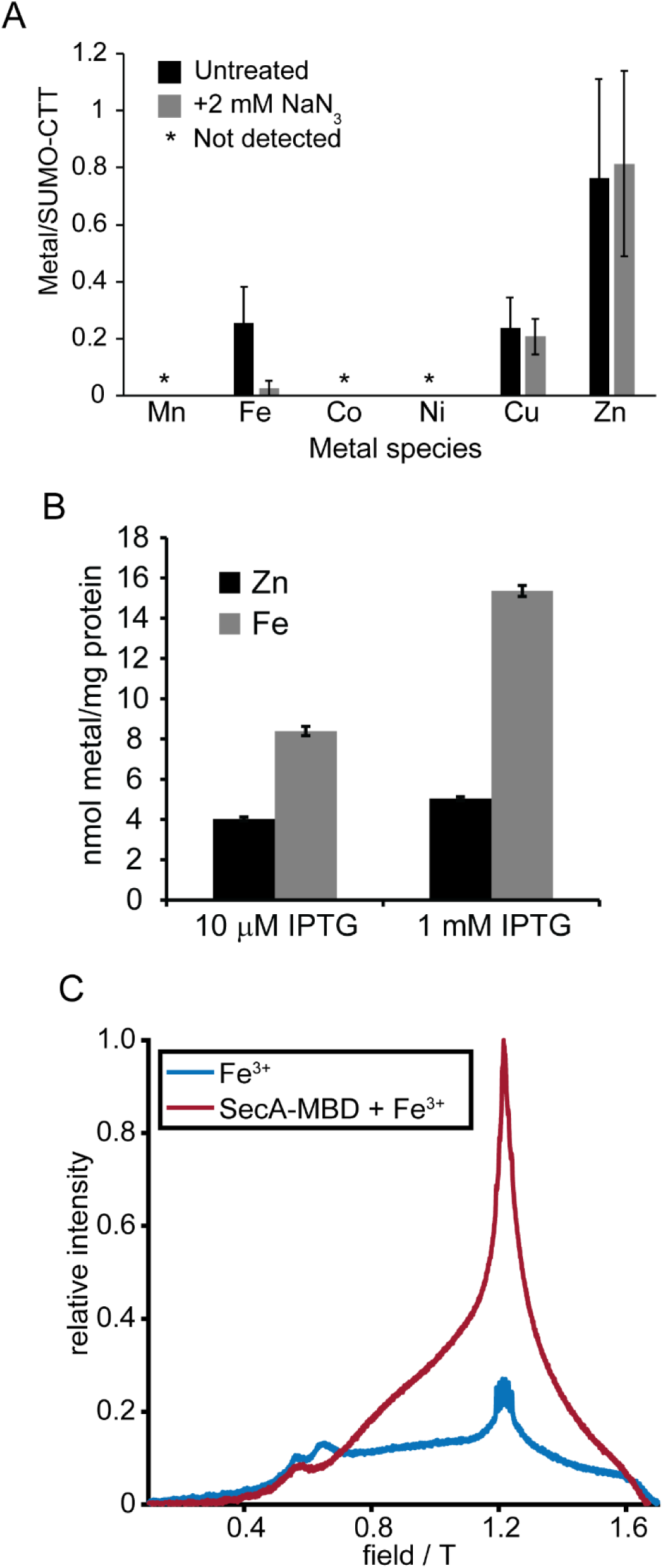
The SecA MBD copurifies with iron and binds to iron *in vitro*. (A) Cells producing SUMO-CTT were incubated in the absence (untreated) or presence (+N_3_) of 2 mM NaN_3_ for 10 minutes. SUMO-CTT was purified from the cell lysates using streptactin beads and washed extensively with buffer. The iron content of SUMO-CTT was determined using mass spectrometry (ICP-MS) and normalised to the protein concentration in the eluted protein. Confidence intervals are one s.d. (B) DRH839 cells (Δ*secA* p_*trc*_-*secA-biotin*) were grown in the presence of 10 μM or 1 mM IPTG to induce expression of SecA-biotin. Cells were rapidly lysed by cell disruption, incubated with streptavidin-sepharose and washed extensively with buffer. The zinc and iron content of the EDTA eluate was determined using ICP-OES and normalised to the amount of protein bound to the streptavidin beads. An estimation of the concentrations of SecA and streptavidin in the purified protein suggested that SecA copurified with stoichiometric amounts of iron. (C) The echo-detected field swept EPR spectrum of oxidised FeSO_4_ was determined in the absence (blue trace) or presence (red trace) of SecA-MBD.

### Co-purification of SecA-biotin with iron

Purification of SecA normally involves overproduction of the protein, followed by multiple purification steps, which could lead to mis-metallation or metal exchange. To determine whether full-length SecA binds to metal *in vivo*, we produced SecA from a chromosomally encoded, IPTG-inducible copy of the *secA* gene. The SecA produced by these strains contained a C-terminal tag that caused it to be biotinylated (SecA-biotin) (30) and allowed us to purify it using streptavidin-coated sepharose beads. SDS-PAGE indicated that the purified protein samples contained only two proteins: SecA and unconjugated streptavidin (**supporting figure S5**). ICP-OES indicated that iron was present at much higher concentrations in the protein samples than zinc (**figure 2B**), suggesting that SecA binds to iron *in vivo*.

### EPR analysis of Fe^3+^ binding by the SecA MBD

Previous work indirectly suggested that the MBD has a lower affinity for iron than for zinc (12). To demonstrate that the MBD of SecA can bind to iron, we investigated the effect of the MBD on the EPR spectrum of Fe^3+^. (Fe^2+^ is not detectable by EPR.) The addition of a metal-free synthetic peptide consisting of the C-terminal 27 amino acids of SecA (SecA-MBD) caused a large increase in the EPR signal of Fe^3+^ at 1.2155 T (∼g = 2) and altered the shape of the EPR spectrum (**figure 2C**). SecA-MBD also inhibited the formation of insoluble precipitates (possibly iron oxides) in solutions containing FeSO_4_ (**supporting figure S6**). These results indicated that SecA-MBD can bind iron.

### Mass spectrometry analysis of metals that co-purify with YecA and YchJ

Fourteen open reading frames in *E. coli* encode polypeptides with the amino acid sequence CXCX_N_CH or CXCX_N_CC: SecA, YecA, YchJ, YjbI, QuuD, ribosomal protein L31, ferredoxin, HCR, YgeH, GlcF, NirB, YjiM, HypA and YeaX (**supporting data S2**). According to UniProt (31), all eight of the proteins with well characterised functions (SecA, L31, ferredoxin, HCR, GlcF, NirB, HypA and YeaX) bind to a metal using all or part of this sequence, and five (ferredoxin, HCR, GlcF, NirB and YeaX) are annotated as binding to iron. Of the remaining proteins, YecA and YchJ contain C-terminal domains, each of which has 73% sequence identity to amino acids 878-899 of SecA. (The C-terminal 22 amino acids of YecA and YchJ are 77% identical to each other.) In addition to the metal-coordinating cysteines, these sequences contain the conserved serine and aromatic residues (**supporting figure S1**). The N-terminal portion of YchJ contains a second CXCX_N_CC motif, which also contains the conserved serine and aromatic residues found in the SecA MBD.

The finding that the SecA MBD binds to iron suggested that YecA and YchJ could also bind to iron. To investigate the metal-binding properties of YecA and YchJ, we purified fusion proteins between YecA or YchJ and hexahistidine-tagged SUMO from *Saccharomyces cerevisiae* and determined the amount of copurifying manganese, iron, cobalt, copper and zinc by ICP-MS. YecA co-purified with zinc at a stoichiometry of 0.84 ± 0.03, consistent with the presence of a single MBD (**figure 3A**). YchJ co-purified with zinc at a stoichiometry of 1.30 ± 0.17 (**figure 3A**), consistent with the presence of two MBDs. Both proteins also copurified with lower, but detectable, amounts of copper (0.17 ± 0.11 for YecA; 0.02 ± 0.01 for YchJ) and iron (0.04 ± 0.03 for YecA; 0.05 ± 0.04 for YchJ).

**Figure 3.**
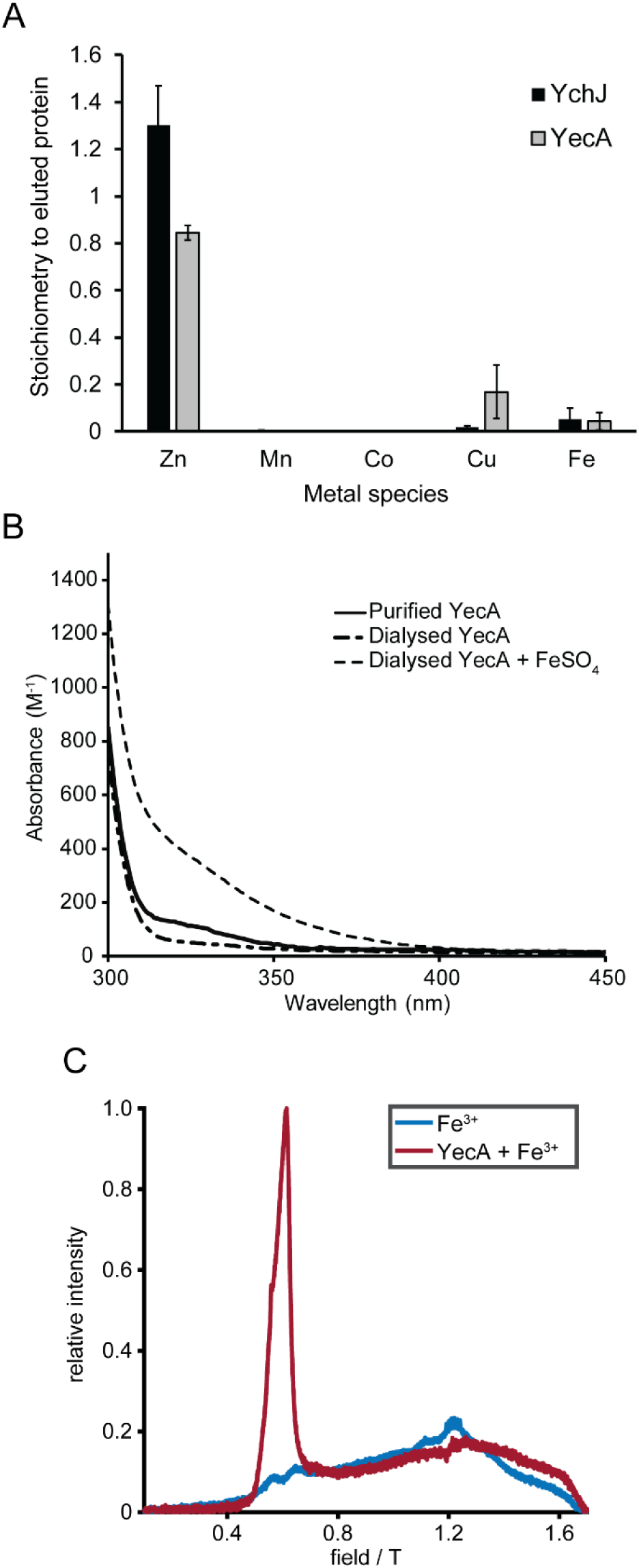
Spectroscopic analysis of metal binding by YecA and YchJ. (A) His-SUMO-YecA and His-SUMO-YchJ were purified using Ni-affinity chromatography, and the amount of co-purifying Zn, Mn, Co, Cu and Fe were determined by ICP-MS. The concentrations of the extracted metals were normalised to the amount of purified protein. Confidence intervals are one standard deviation. (B) Buffer subtracted absorbance spectra of purified YecA (solid black), purified YecA that was dialysed against EDTA (dash-dot) and EDTA-dialysed YecA to which an equimolar concentration of FeSO_4_ was added (dashed). The absorbance spectra were normalised to the concentration of YecA. (C) The echo-detected field swept EPR spectrum of FeCl_3_ was determined in the absence (blue trace) or presence (red trace) of YecA.

### Absorbance spectroscopy of purified YecA and YchJ

During purification, the colour of the YecA- and YchJ-bound Ni-NTA columns was yellow. If TCEP was included in the buffers used for purification of YecA, the Ni-NTA column remained yellow after elution of the bound protein. Ni-NTA columns are blue-green due to the coordinated Ni^2+^ ion and normally regain this colour after elution of the bound protein with imidazole. The change in colour suggested that the bound Ni^2+^ had exchanged with another metal ion, presumably from the overexpressed protein in the cell lysate used for purification. Analysis of the EDTA eluate indicated that the bound metal ion was iron (**supporting figure S7**). Furthermore, purified YecA and YchJ were pale yellow in colour (Zn-bound proteins are normally colourless) and gradually became colourless with incubation at 4°C—typically over a period of one to several hours. YecA absorbed light with a peak at ∼330 nm, and YchJ absorbed light with a peak at ∼340 nm (**supporting figure S8**). Removal of the N-terminal His-tagged SUMO did not affect the absorbance spectrum of YecA. Dialysis of YecA against buffer containing EDTA resulted in a decrease in absorbance at 330 nm (**figure 3B**), and the addition of an equimolar concentration of FeSO_4_ restored absorbance of YecA at 330 nm (**figure 3B**). Taken together, these results suggested that the yellow colour of the purified YecA protein was due to coordination of Fe^2+^ (or potentially Fe^3+^), and the slow conversion of the purified protein to colourless suggested that binding to iron was labile under the experimental conditions. Dialysis of YchJ against EDTA resulted in aggregation. Consequently, we did not investigate YchJ further.

### EPR analysis of Fe^3+^ binding by YecA

To confirm that YecA can bind iron, we investigated the interaction of YecA with Fe^3+^ using EPR spectroscopy. The addition of a stoichiometric amount of purified metal-free YecA to a solution of Fe^3+^ (**figure 3C, blue line**) resulted in a new feature at 0.6145 T that is ∼g = 4 (**figure 3C, red line**), indicating that there was a change in the electronic environment of the iron and indicating that YecA can bind to Fe^3+^.

### Identification of the iron-coordinating domain in YecA by NMR spectroscopy

To determine which domain was responsible for binding to iron, we purified ^15^N- and ^13^C-labelled YecA and investigated its structure using NMR spectroscopy. We could assign the resonances for the C-terminal 20 amino acids in the TROSY spectrum of the protein (excluding Pro-206 and Pro-208, which do not contain an N-H bond) (**figure 4A**). Several amino acids, including Arg-203, Asp-204, Asp-205, Leu-220 and His-221, produced two resonances, suggesting that the MBD in the metal-free protein exists in two distinct conformations (**figure 4A**). The addition of FeSO_4_ resulted in the broadening and flattening of most of the resonances from amino acids in the MBD due to the paramagnetic properties of iron, but it affected very few of the resonances from the N-terminal ∼200 amino acids (**figure 4B**). This result suggested that the MBD is responsible for the iron-binding activity of YecA.

**Figure 4.**
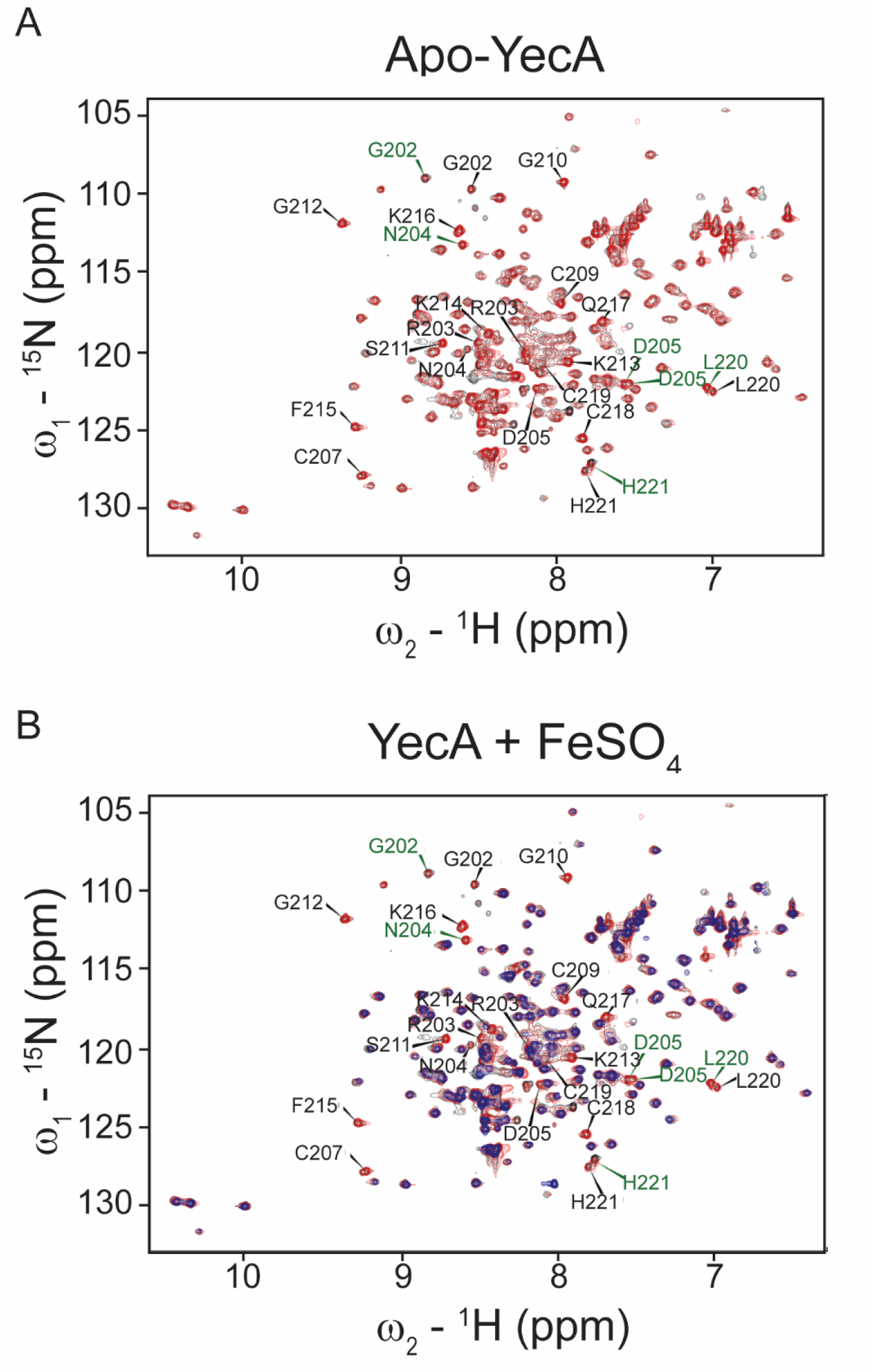
YecA binds to iron *via* its MBD. (A) TROSY spectrum of metal-free ^15^N & ^13^C-labelled YecA. The ^1^H, ^15^N, and ^13^C resonances of the polypeptide backbone for the C-terminal 20 amino acids (excluding prolines-206 and -208) were assigned using triple resonance experiments. Resonances of amino acids in the MBD are indicated with arrowheads. Amino acids producing two resonance peaks are labelled in green. (B) Overlay of the TROSY spectra of metal-free YecA (red) and YecA in the presence of equimolar concentrations of FeSO_4_ (blue). The absence of a blue peak suggests that the amino acid producing the resonance is in the proximity of the bound iron.

### ^1^H-NMR analysis of metal binding by SecA-MBD

To investigate the relative affinity of the SecA MBD for zinc and iron, we examined binding of the SecA-MBD peptide to iron and zinc using ^1^H-NMR. The addition of ZnSO_4_ to SecA-MBD resulted in multiple changes in its ^1^H-NMR spectrum including the appearance of resonances in the 8.5-9.5 ppm region, which are indicative of the formation of secondary structure (**figure 5A, blue and green traces**), consistent with previous studies (13). As with YecA, the addition of FeSO_4_ resulted in a broadening and flattening of most of the ^1^H resonances due to the paramagnetic properties of iron (**figure 5A, red trace**). The reduction in signal was more pronounced for ^1^Hs that were nearer to the predicted metal binding site compared to those further away from the metal-binding site. FeSO_4_ did not affect the spectra of control peptides that do not bind iron, indicating that the change in signal was caused by binding to iron.

**Figure 5.**
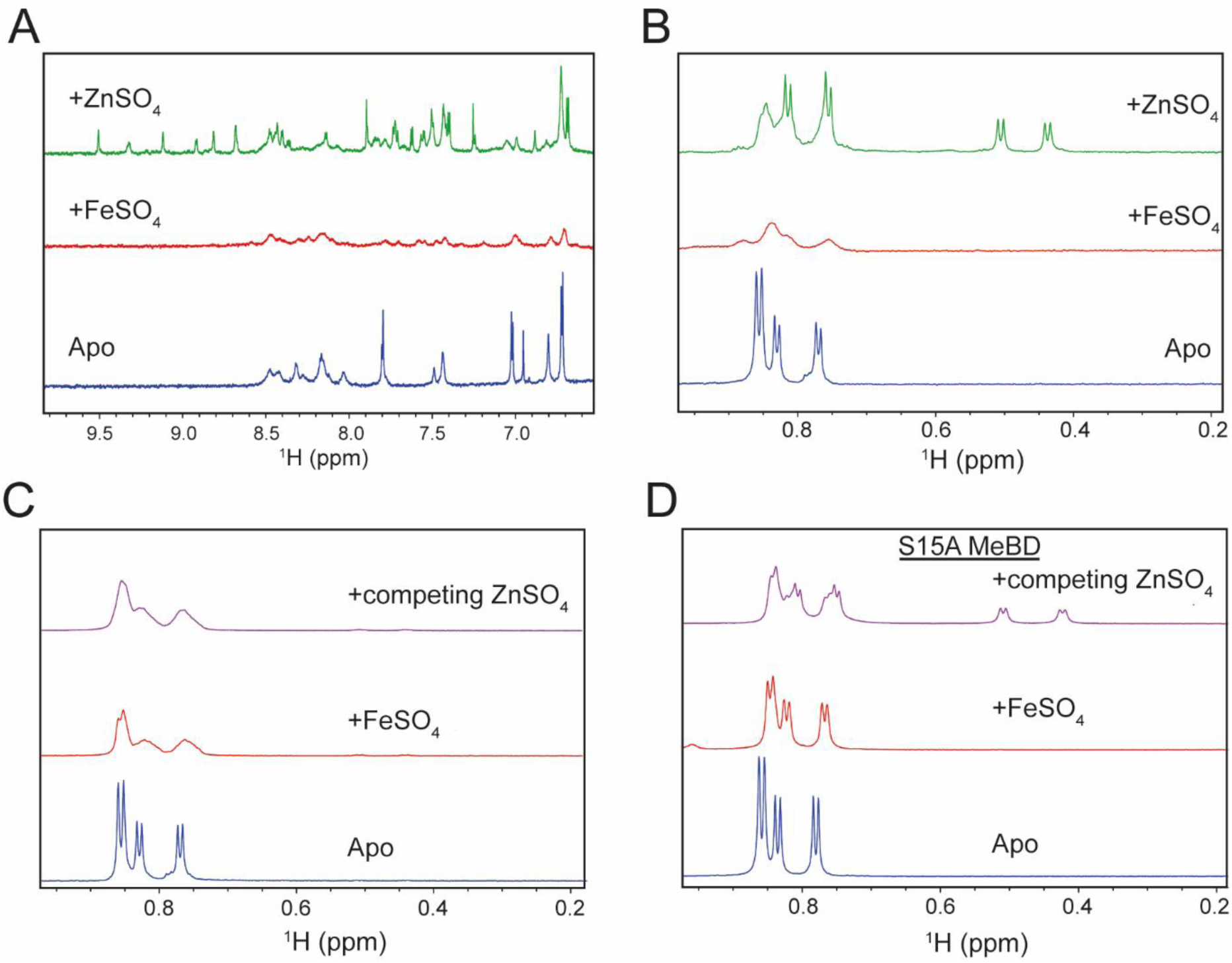
^1^H-NMR analysis of metal binding by SecA-MBD and SecA-MBD^S889A^. (A) 6.5-10 ppm region of the ^1^H-NMR spectra of SecA-MBD in the absence (blue) or presence of equimolar concentrations of FeSO_4_ (red) or ZnSO_4_ (green). The appearance of resonances in the ∼8.5-10 ppm region in the presence of ZnSO_4_ indicates the formation of secondary structure. The strong quenching of the resonances in the presence of FeSO_4_ is due to proximity to the bound iron ion. (B) 0.2-1.0 ppm region of the ^1^H-NMR spectra of SecA-MBD in the absence (blue) or presence of equimolar concentrations of FeSO_4_ (red) or ZnSO_4_ (green). Binding of SecA-MBD to zinc results in a shift of the resonances corresponding to the methyl ^1^Hs of valine, and binding of SecA-MBD to iron results in a significant broadening and flattening of these resonances. (C & D) 0.2-1.0 ppm region of the ^1^H-NMR spectra of SecA-MBD (C) or SecA-MBD^S889A^ (D) in the absence of metal (blue), in the presence of equimolar FeSO_4_ (red) and after the addition of competing concentrations of ZnSO_4_ to the iron-bound peptide after >10 minutes of equilibration at room temperature (purple).

We next investigated the ability of zinc to compete with iron for binding to SecA-MBD. To this end, we monitored the ^1^H resonances from the valine methyl groups, which displayed differences that were distinctive for the apo-, Zn-bound and Fe-bound states (**figure 5B**). Previous work indicated that the dissociation rate of Zn^2+^ from the MBD is relatively rapid (half-life of minutes) (12). However, the addition of ZnSO_4_ to SecA-MBD that had been pre-incubated with FeSO_4_ did not cause a detectable change in the ^1^H-NMR spectrum, even after ∼40 minutes of incubation (**figure 5C**), suggesting that SecA-MBD binds preferentially to iron.

### Role for a conserved serine in metal preference

The SecA MBD contains a nearly invariant serine residue, Ser-889 (**supporting figure S1**) (10,13), which is conserved in the MBD of YecA and both MBDs of YchJ (**supporting figure S9**). Previously published structural models of the SecA MBD indicate that the side chain of Ser-889 points inward toward the metal ligand and is not directly involved in binding to SecB (or likely ribosomes) (13,14,32) (**supporting figure S10**). In addition, the N-terminal MBD of YchJ, which lacks the amino acid residues required for binding to SecB, contains a serine at the same position suggesting that the serine is required for the structure of the MBD. In support of this notion, ^1^H-NMR analysis indicated that SecA-MBD containing an alanine substitution at this position (SecA-MBD^S889A^) exchanged iron for zinc at a detectable rate (**figure 5D**). These results suggested that Ser-889 is important for the folding of the MBD and/or coordination of the metal ion.

### Binding affinity of SecA-MBD peptides for Zn^2+^

To determine whether the alanine substitution caused a general defect in metal binding, we determined the affinity of SecA-MBD and SecA-MBD^S889A^ for Zn^2+^ using ITC. The affinity of the wild-type SecA-MBD for Zn^2+^ (36.0 ± 11.6 nM; **supporting figure S11A**) was not significantly different from that of SecA-MBD^S889A^ (46.0 ± 9.0 nM; **supporting figure S11B**), suggesting that the substitution does not affect its affinity for Zn^2+^. We could not determine the affinity for SecA-MBD for iron due to the interfering heat exchange caused by aerobic oxidation of Fe^2+^ and because binding of SecA-MBD to Fe^3+^ did not cause a detectable heat exchange. Nonetheless, these results, taken together with the ^1^H-NMR experiments, suggested that the serine-to-alanine substitution specifically affects the affinity of SecA-MBD^S889A^ for iron.

### Modelling octahedral coordination of a metal by the SecA MBD

Because the strong reduction in signal in the ^1^H-NMR spectrum of SecA-MBD caused by binding to iron made it difficult to determine the structure of the SecA MBD bound to iron directly, we modelled the structure of the MBD bound to iron using molecular dynamics. Fe^2+^ and Fe^3+^ can be hexavalent and are frequently coordinated in an octahedral geometry (33). We therefore simulated the MBD coordinated octahedrally to a hexavalent metal ion. In addition, we assumed that the hydroxyl group of Ser-889 and a hydroxyl ion from the solvent bound the two additional coordination sites of the metal. Finally, because it was not clear whether the δ- or ε-nitrogen of His-897 binds the metal, we simulated both possibilities. These simulations suggested that the imidazole ring of His-897 is out of plane with the metal ion (**figure 6**), and longer simulations using reduced constraints on the metal-coordinating atoms suggested that binding *via* the ε-nitrogen (**supporting figure S12A**) was more stable than *via* the δ-nitrogen (**supporting figure S12B**), consistent with the structures of other iron-binding proteins (34). Tetrahedral coordination of zinc was previously reported to induce strain into the polypeptide backbone (13). However, Ramachandran plots of the structures generated by our simulations indicated that the MBD can coordinate iron octahedrally without inducing stress in the polypeptide backbone (**supporting data S3 and S4**). The SecB-binding surface was not significantly perturbed in the iron-bound structural models compared to previously published structures (**supporting data S3 and S4**) (13,14). These results indicate that Ser-889 can participate in coordination of a hexavalent metal, such as iron.

**Figure 6.**
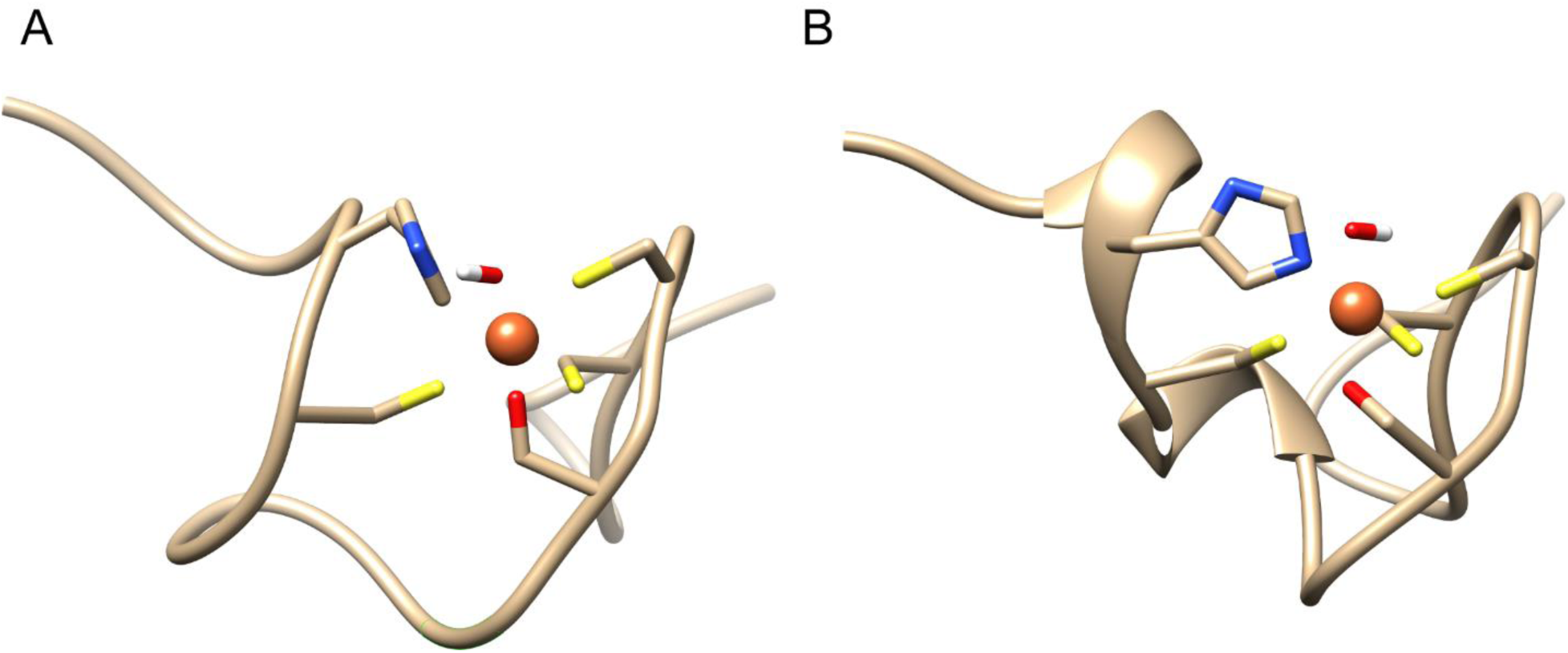
Structural models of octahedral metal coordination. Ribbon diagrams of example structural models of the MBD coordinating a hexavalent metal using the δ-nitrogen (A) and the ε-nitrogen (B) of His-897 generated using molecular dynamic simulations. Example structures from each 1 ns of the final 10 ns of each 100 ns simulation can be found in the online supporting data. The side chains of the metal-coordinating amino acids are depicted as sticks (yellow, sulfur; blue, nitrogen; red, oxygen). Images were rendered using UCSF Chimera v. 1.12 (72).

## DISCUSSION

Our results indicate that iron is a physiological ligand of SecA-like MBDs. The SecA CTT, YecA and YchJ copurify with significant amounts of iron, and full-length SecA copurifies predominantly with iron when rapidly purified from cells producing it at low levels. Biophysical experiments indicate that the MBDs of SecA and YecA can bind to iron, and competition experiments suggest that the MBD of SecA binds preferentially to iron. Finally, our results indicate that azide treatment causes autoinhibition of SecA *in vivo* by disrupting the interaction of the MBD with iron. Taken together, our results suggest that a significant proportion of SecA is bound to iron *in vivo* and that the iron-bound form could play a physiological role in regulating the activity of the protein.

Our results do not exclude the possibility that both iron and zinc are physiological ligands of the MBD. For example, it is possible that the relative abundances of the zinc- and iron-bound forms of SecA are involved in autoregulation of SecA by the CTT. Indeed, the Zn-bound form of the protein appears to be functional *in vitro* (12,14,35,36). However, the known abundances of zinc and iron in the cell suggest that the iron-bound form of SecA is the dominant species *in vivo*. Although the affinity of the SecA-MBD-iron complex could not be determined directly, the preference of the MBD for iron over zinc suggests that the K_D_ is less than 40 nM. The cytoplasmic concentration of iron is normally ∼50 nM while the concentration of free zinc is fM to pM (33,37,38). In addition, although metal-binding proteins bind promiscuously to different transition metals, they typically bind with higher affinity to zinc than to iron (33,38), making the higher affinity of the SecA MBD for iron remarkable. These results suggest that the MBD is normally bound to iron *in vivo*. The strong conservation of Ser-889 suggests that iron binding is evolutionarily conserved.

Our molecular dynamics simulations suggest that the specificity of the MBD for iron could be determined by the binding geometry. Zinc is nearly always bound using a tetrahedral geometry while iron can be coordinated using several geometries, including octahedral (33,38). Previous work has established that Cys-885, Cys-887, Cys-896 and His-897 are required for binding to metal (12,14). Our experimental results suggest the conserved serine also participates in coordination of the bound metal ion, suggesting that the bound metal ion is coordinated using at least five sites. It is not yet clear if the bound iron is coordinated at a sixth site and if so, which amino acid binds the iron. Our simulations used a solvent molecule to coordinate the bound metal, but it is possible that the conserved aromatic residue at position 893 indirectly participates in coordination of the metal (13). For example, the π-ring system of Tyr-893 could coordinate a solvent molecule that participates in metal-ion coordination (39). Although π-electron systems can coordinate iron (40), preliminary simulations suggest that binding of Tyr-893 to the metal results in steric clashes with the other metal-coordinating side-chains.

Our results provide a potential explanation for the discrepancy between the concentrations of azide required to inhibit SecA *in vivo* and *in vitro*. Previous work indicates that azide binds to the ADP-bound form of SecA and inhibits nucleotide exchange (19,20). Our results suggest that azide additionally disrupts the structure of the iron-bound MBD, resulting in autoinhibition of SecA. Our results suggest that SecA detaches from the cytoplasmic membrane as a consequence of disrupting the MBD. Previous research indicates that CTT-mediated autoinhibition is not sufficient to completely inhibit growth of *E. coli* (10,35). However, our results suggest that autoinhibition slows bacterial growth at subinhibitory concentrations of azide, which likely enhances the inhibitory effect of azide on the ATPase activity of SecA at inhibitory concentrations. We cannot rule out the possibility that the role of the bound iron ion is purely structural. Indeed, many helicases contain evolutionarily conserved iron-sulfur clusters that appear to function solely as structural scaffolds (41). However, our results raise the possibility that the MBD senses an environmental stress and that azide treatment mimics this stress. For example, the MBD could sense changes in the oxidation state of the bound iron (Fe^2+^ versus Fe^3+^), the identity of the bound metal (Zn versus Fe) or the oxidation state of the metal coordinating cysteines (42-44).

The functions of YecA and YchJ are unknown. Both proteins contain domains of unknown function (UPF0149 in YecA and UPF0225 in YchJ (45)) and are found in a broad range of bacterial species. Neither protein is universally conserved (31) or essential for viability (23,46). However, the conservation of the amino acids that mediate the interaction between the SecA MBD and SecB in the C-terminal MBDs of YecA and YchJ suggest that that these proteins could also potentially interact with the SecB or ribosomes. Large scale genetic screens do not suggest an obvious function for either proteins (46,47), suggesting that YecA and YchJ do not carry out their functions under laboratory growth conditions or are not normally produced (or both). The sole publication mentioning *yecA* (48) suggests that it could encode the gene *ssaG* (*ssa* mutants suppress the temperature sensitive growth defect caused by a *secA51* mutation) (49). However, whole genome sequencing of strain DO314 (which contains an *ssaG1* mutation) suggests that the suppressor mutation does not map to the *yecA* gene (unpublished result).

The evolutionary conservation of the MBD supports the idea that YecA and YchJ interact with the ribosome. In addition to recent work suggesting that MBD of SecA binds to the ribosome (10), the methionine amino peptidases of many bacterial species contain a SecA-like MBD (45). (In *E. coli*, methionine amino peptidase binds to the ribosome (50).) In addition, ribosomal protein L35a in *Drosophila melanogaster* contains a SecA-like MBD, suggesting that this role is evolutionarily conserved. Our results suggest that SecA-like MBDs could also regulate interactions with the ribosome in response to environmental stress.

## EXPERIMENTAL PROCEDURES

### Chemicals and media

All chemicals were purchased from Fisher or Sigma-Aldrich unless indicated. Synthetic peptides were synthesised by Severn Biotech (Kidderminster, UK) or using an in-house synthesiser. The quality of the peptides was checked using MALDI mass spectrometry. 100X EDTA-free protease inhibitor cocktail was purchased from Pierce (Thermo-Fisher). Cells were grown using LB medium (51). Where indicated, IPTG was added to the culture medium. Where required, kanamycin (30 μg/ml) was added to the growth medium.

### Strains and plasmids

Strains and plasmids were constructed using common genetic methods (51). Single gene deletion mutants from the Keio collection (46) were obtained from the *E. coli* genetic stock centre (CGSC; Yale University, New Haven, Connecticut). Strain MG1115 (52) was a kind gift from M. Grabowicz and T. Silhavy. Strain DO314 (which contains an *ssaG1* mutation) was a kind gift from D. Oliver. All other strains were lab stocks.

### Azide sensitivity assays

Filter disc assays were conducted as described by Huie and Silhavy (17) except that bacterial lawns were produced by evenly swabbing an overnight culture of *E. coli* over the surface of an LB containing the indicated concentrations of EDTA, ZnSO_4_ or FeSO_4_. An empty 6 mm antibiotic assay filter disc (Merck-Millipore) was then placed in the centre of the lawn and 10 μl of 1 M NaN_3_ was pipetted onto the filter disc. The lawns were then grown overnight at 37°C.

### TraDIS

TraDIS experiments were conducted as described previously (22,23). 50 ml of LB broth containing 0, 0.25 or 0.5 mM NaN_3_ were inoculated with 10 μl of a library of ∼1 million *E. coli* BW25113 mini-Tn*5* insertion mutants, and the cultures were grown to OD_600_ 1.0. Genomic DNA was extracted using a Qiagen QIAamp DNA blood mini kit and then processed using a two-step PCR method (53), which results in Illumina-compatible products. The PCR products were purified using the Agencourt AMPure XP system by Beckman Coulter. The products were sequenced using an Illumina MiSeq sequencer and the reads were mapped to the *E. coli* reference genome NC_007779.1 (*E. coli* K-12 substr. W3110). The number of insertions in the coding sequences (CDS) for each gene was then determined. To reduce the number of false positives due to sequencing assignment errors, genes that exist in multiple copies on the chromosome (e.g. insertion elements and rRNA operons) were eliminated from the analysis. Genes containing 15 or fewer total insertions across all three conditions were eliminated in the data presented in **figure 1A**.

### Phospholipid binding assay

Strains producing His-SUMO-SecA (DRH625; (53)) or His-SUMO-SecA^C885A/C887A^ (MJ118; (10)) were grown to late log phase in LB and induced using 1 mM IPTG. For *in vivo* experiments, the culture was divided after the addition of IPTG, and half of the culture was treated with 2 mM sodium azide for 10 minutes. Cells were harvested by centrifugation, resuspended in lysis buffer (20 mM HEPES [potassium salt] pH 7.5, 500 mM NaCl, 1 mM tris(2-carboxyethyl)phosphine [TCEP]) and lysed by cell disruption. Clarified cell lysates were passed over a 1 ml Ni-HiTrap column (GE Biosciences), and the bound protein was washed with 50 volumes of lysis buffer containing 20 mM Imidazole. His-SUMO-SecA was eluted from the column using lysis buffer containing 500 mM imidazole and dialysed against wash buffer lacking imidazole. The concentration of the protein was adjusted to 1 mg/ml to a final volume of 2 ml, and the phospholipids were extracted using 2 ml methanol and 1 ml chloroform. The chloroform phase was removed, and phospholipids were concentrated by air drying and resuspending in chloroform to a final volume of 100 μl. The indicated volume of sample was spotted onto TLC silica gel 60 T_254_ plates (Merck Millipore) and resolved using a mixture of 60 chloroform: 25 methanol: 4 water. For the *in vitro* phospholipid release assay, 0.2 mg of His-SUMO-SecA purified from untreated cells was bound to 100 μl Ni-NTA agarose beads (Life Technologies). The beads were incubated with 500 μl of lysis buffer or buffer containing 2 mM sodium azide for 10 minutes and washed three times with 500 μl lysis buffer. The protein was eluted from the beads with lysis buffer containing 500 mM imidazole, and the phosopholipid content of the supernatant was analysed using the method described above.

### Determination of metal content of Strep-SUMO-CTT

To determine the effect of azide on binding of Strep-SUMO-CTT to iron, 100 ml cultures of BL21(DE3) containing plasmid pDH543 (10) were grown to OD_600_ 1.0 in LB, and production of Strep-SUMO-CTE was induced using 1 mM IPTG. After 1 hour, cultures were split, and half the culture was treated with 2 mM NaN_3_ for 10 minutes. Cells were rapidly cooled, harvested by centrifugation and lysed using B-PER cell lysis reagent from Pierce (Rockford, Illinois). Cell lysates were incubated for 15 minutes with 50 μl of a 50% Streptactin-Sepharose slurry that had been pre-equilibrated with wash buffer (10 mM HEPES (potassium salt) pH 7.5, 100 mM potassium acetate) and washed extensively with wash buffer. The samples were washed a final time using 10 mM HEPES (potassium salt) pH 7.5 to remove excess salt, and the total protein was eluted off of the column using using 10 mM HEPES (potassium salt) pH 7.5 buffer containing 7M guanidinium hydrochloride. The propensity to aggregate was determined by diluting 50 μl of the guanidinium-denatured protein into 950 μl of a 20 mM HEPES buffer and measuring light scattering at 500 nm. The metal content of the samples was determined using ICP-MS (School of Geography, Earth and Environmental Sciences, University of Birmingham, UK).

### Determination of metal content of SecA-biotin

100 ml cultures of DRH839 (MC4100 Δ*secA* λ-p_*trc*_-*secA-biotin*..SpecR) (30) were grown in 10 μM or 100 μM IPTG to OD_600_ ∼ 1. Cells were lysed using cell disruption, and lysates were incubated with 100 μl Streptactin-sepharose (IBA Lifesciences, Göttingen, Germany) for 15 minutes. The beads were washed four times with 30 ml buffer (50 mM potassium HEPES, pH 7.5, 100 mM potassium acetate, 10 mM magnesium acetate, 0.1% nonidet P40). Metal was eluted from the beads by incubating with 10 mM HEPES, pH 7.5, 50 mM EDTA) at 55°C for 30 minutes, and the zinc and iron content was determined using ICP-OES. The amount of bound protein was determined by boiling in SDS sample buffer and analysing using Bradford reagent (BioRad, Hercules, CA). The eluted protein was resolved on a BioRad 15% TGX gel. The metal content was determined using ICP-OES (School of Geography, Earth and Environmental Sciences, University of Birmingham, UK).

### Purification of YecA and YchJ

The *yecA* and *ychJ* genes from *Escherichia coli* K-12 were fused in-frame to the 3′ end of the gene encoding SUMO from *S. cerevisiae* in plasmid pCA528 and purified as described previously (54). For identification of the copurifying metals and absorbance spectroscopy, BL21(DE3) cells containing the SUMO-YecA and SUMO-YchJ plasmids were grown to OD_600_ 1.0 at 37°C. Cells were then shifted to 25°C and grown overnight in the presence of 1 mM IPTG. Cells were lysed in buffer 1 (20 mM potassium HEPES, pH 7.5, 100 mM potassium acetate, 10 mM magnesium acetate) containing protease inhibitor by cell disruption.

When noted, TCEP was added to buffer 1 during lysis at a concentration of 1 mM. Lysates were passed over a 1 ml His-Trap HF column (GE Healthcare). The bound protein was washed with 15 ml high salt wash buffer (20 mM potassium HEPES, pH 7.5, 500 mM potassium acetate, 10 mM magnesium acetate, 50 mM imidazole and 15 ml low salt wash buffer (20 mM potassium HEPES, pH 7.5, 100 mM potassium acetate, 10 mM magnesium acetate, 50 mM imidazole). The bound protein was eluted using elution buffer (20 mM potassium HEPES, pH 7.5, 100 mM potassium acetate, 10 mM magnesium acetate, 500 mM imidazole). The eluted protein was dialysed against buffer 1 to remove the imidazole and concentrated using concentrators with a 5 kDa cutoff (Vivaspin).

For HSQC-NMR analysis of YecA, cells containing the YecA plasmid were grown in M9 minimal media containing ^15^NH_4_Cl, ^1^H_7_-^13^C_6_-glucose (2 g/l) and trace minerals. Cultures were grown to OD_600_ 0.8 at 37°C, shifted to 18°C and induced overnight with 1 mM IPTG. Cells were lysed by cell disruption and purified using a 1 ml His-Trap column as described above. After elution, the protein was treated with purified hexahistidine-tagged Ulp1 from *S. cerevisiae* to remove the SUMO tag and dialysed overnight against buffer 2 (20 mM potassium HEPES, pH 7.5, 100 mM potassium acetate, 10 mM magnesium acetate, 5 mM β-mercaptoethanol) to remove the imidazole. The SUMO tag and Ulp1 protease were removed from the purified protein by passing the cut protein over a 1 ml His-Trap column. Purified YecA was then concentrated using anion exchange chromatography, and the eluate was dialyzed against buffer 3 (20 mM potassium HEPES, pH 7.5, 100 mM potassium acetate, 10 mM magnesium acetate, 5 mM β-mercaptoethanol, 1 mM EDTA) and finally against buffer 1 to remove the β-mercaptoethanol and EDTA.

### Metal ion analysis

The metal ion content of purified YecA and YchJ was determined using ICP-MS (School of GEES, University of Birmingham). The 5 kDa MWCO concentrator filtrate (Sartorius, Göttingen, Germany) was used to control for the amount of unbound metal in the protein samples. The zinc and iron ion content of the EDTA eluate from the Ni-NTA column after purification in the presence of 1 mM TCEP was determined using ICP-OES (School of Geography, Earth and Environmental Sciences, University of Birmingham).

### Absorbance spectroscopy

The absorbance spectra of 200 μl of 600-800 μM purified YecA or YchJ in buffer 1 were determined from 300-600 nm using a CLARIOstar plate reader (BMG Labtech) using UV-clear flat bottomed 96-well plates (Greiner). The absorbance spectrum for the buffer alone was subtracted from that of the purified protein, and the absorbance was normalised to the concentration of the protein in the sample.

### EPR spectroscopy

EPR samples were suspended in 50 μl buffer 1 containing 30% glycerol. For YecA, samples contained 0.85 mM FeCl_3_ or 0.6 mM YecA and FeCl_3_. For SecA-MBD, samples contained 0.5 mM FeSO_4_, which had been left to oxidize aerobically in a 25 mM water/glycerol stock, with 0.5 mM SecA-MBD or just the metal salt in buffer. Measurements were taken with a Bruker Elexsys E580 spectrometer with an ER 5106QT-2w cylindrical resonator operating at 34 GHz, i.e. Q-band. Quartz tubes with 3 mm O.D. were used. Experiments were conducted at 10 K using a cryogen free variable temperature cryostat (from Cryogenic Limited). Echo-detected field sweeps were conducted by sweeping from 0.1 to 1.7 T with 4000 points using a Hahn echo sequence where the π pulse length was 32 ns and the time between pulses was 400 ns. The power level was determined by observing the maximum echo and was around 0.38 mW which is indicative of a high-spin system. The shot repetition time was set at 100 μs with 50 shots per point which was sufficient for the iron, though caused some saturation of the Mn^2+^ contaminant peak. The resultant echo-detected field swept profiles were normalised for plotting, taking account of differences in video gain, concentration and numbers of averages.

### NMR backbone assignment of YecA

Assignment spectra were acquired on 0.5 mM ^15^N, ^13^C-labelled protein in 20 mM [MES]-NaOH [pH 6.0], 10 mM NaCl, 10% D_2_O in a 5 mm Shigemi tube (Shigemi Inc.). The ^1^H, ^15^N, and ^13^C resonances of the YecA backbone were assigned using BEST TROSY versions of HNCA, HN(CO)CA, HNCACB, HN(CO)CACB, HNCO and HN(CA)CO (55-61). All experiments were performed at 298 K and acquired with a spectral width of 14 ppm in ^1^H, collecting 1024 real data points, and 30 ppm in ^15^N, collecting 92 increments using a Bruker 900 MHz spectrometer equipped with a 4-channel AVANCE III HD console and a 5 mm TCI z-PFG cryogenic probe The centre of the spectra was set to 4.698 ppm in the ^1^H and 118 ppm in ^15^N. All spectra were acquired collecting 128 increments in the ^13^C dimension using a non-uniform sampling scheme. The HN(CO)CACB and HNCACB experiments were acquired using 64 scans per increment, a spectral width of 76 ppm in the ^13^C direction with the centre around 43.665 ppm. The HNCA and the HN(CO)CA experiments were acquired using 32 scans per increment, a ^13^C spectral width of 30 ppm with the centre of the spectra set to 55.9 ppm. The HN(CA)CO and HNCO experiments were acquired using 32 scans per increment, a ^13^C spectral width of 16 ppm centred around 176.2 ppm. Non-uniformed sampled data were reconstructed using the compressed sensing algorithm with MDDNMR (62) and processed using nmrPipe (63). Spectra were analysed in Sparky (64).

### ^1^H-NMR spectroscopy

All spectra were obtained at 298 K on a Bruker 900 MHz spectrometer equipped with a cryogenically cooled 5 mm TCI probe using excitation sculpting for water suppression on a sample in 90% H_2_O/10% D_2_O. Sequence specific assignments were completed using a TOCSY experiment in 90% H_2_O/10% D_2_O using a DIPSI2 spin-lock with a mixing time of 65 ms, 32 transients and collecting 512 increments with a spectral width of 10 ppm in both dimensions. 1D data sets comprised 16 transients, 32000 data points and a spectral width of 16 ppm. All data were processed using Topspin 3.2.6 software using an exponential window function with a line broadening of 1 Hz.

### ITC

ITC measurements were conducted in a MicroCal VP-ITC calorimeter (Piscataway, NJ, USA). All solutions were centrifuged for 5 min at 13,000 rpm and then thoroughly degassed under vacuum for 5 min with gentle stirring immediately before use. 0.1 mM ZnSO_4_ was titrated into a solution of the indicated peptide (0.01 mM) in the sample cell (Vo = 1.4037 ml). Titrations consisted of a preliminary 2 µl injection followed by 50 6 µl injections of 12 s duration with 700 s between each injection. All experiments were conducted at 25°C with an initial reference power of 10 µcal/s. The raw data were analysed with Origin 7.0 using one-binding site model and were corrected for the heat of dilution of the metal ion in the absence of peptide.

### Molecular dynamics simulations

Energy minimisations and molecular dynamics simulations were performed using Amber 18 (65). All parameters were from the Amber force field ff14SB (66), except where noted below. Charges for the deprotonated serine (renamed SEO) were determined by comparison with deprotonated cysteine (Amber residue name CYM) (**supporting table S2)**. Parameters for the Fe^2+^ bound histidine (His 897) and HO-were as previously described (67). An SPCE water model and a 2+ iron ion were used, as previously described (68-71). The C-terminal 24 amino acids of SecA (KVGRNDPCPCGSGKKYKQCHGRLQ) were used for modelling. Initial molecular coordinates were derived from the superposition of structures 1SX1 (13) (residues 1 to 21) and 1TM6 (32), followed by energy minimisation, to remove any strain arising from the fusing of the two coordinate sets. The system was then solvated in an octahedral box of water extending 8 Å from the protein surface. Ewald summation was applied to the simulations with a non-bonded cut-off of 10 Å. The system was minimised with positional restraints (500 kcal/(mol•Å)) on every atom of the peptide and on the iron. This was followed by further minimisation without positional restraints but with weak restraints to enforce an octahedral geometry. Distance restraints were applied between the iron and the six chelating atoms (10 kcal/(mol•Å)), as indicated in the legend of **figure S7**, and between atoms on adjacent vertices of the octahedron, listed in **supporting table S4** (2.5 kcal/(mol•Å)). A torsion angle restraint (10 kcal/(mol•rad)) was applied to the sulfur atoms of the residues Cys885, Cys887 and Cys896 and to the δ or ε nitrogens of His897. **Supporting table S3** presents a list of all the restraint parameters. The system was then heated to 300 K in six 50 K steps, using an NVT ensemble and a Langevin thermostat with *γ* = 1 ps^−1^. The peptide and iron were subjected to positional restraints (10 kcal/(mol•Å)), but no other restraints, during the entire heating process. Subsequently, the system was equilibrated for 1.2 ns using an NPT ensemble and a Langevin thermostat with *γ* = 1 ps^−1^. Residues directly involved in the octahedral coordination of iron had a constant positional restraint of 10 kcal/(mol•Å) applied for 600 ps, followed by application of the distance and torsional restraints previously defined. Residues not directly chelated to the iron also had positional restraints initially of 10 kcal/(mol•Å), but reduced to 1 kcal/(mol•Å) at 200ps, 0.1 kcal/(mol•Å)) at 400 ps and zero from 600 ps onwards. A 100 ns long production simulation was performed in an NPT ensemble with a Berendsen thermostat (*τ* = 10 ps). The same distance and torsional restraints were applied as described for the minimisation. This 100 ns simulation was then extended for a further 20 ns, for a total of 120 ns, but with the force constants reduced from 10 to 2 kcal/(mol•Å), 2.5 to 0.5 kcal/(mol•Å) and 10 to 2 kcal/(mol•rad) for distances between iron and chelating atoms, distances between adjacent atoms on the surface of the octahedron and torsion angle, respectively. The above protocol was followed once with the restraints between His-897 and iron via the δ-N and once with the restraints via the ε-N.

## DATA AVAILABILITY STATEMENT

HSQC and ^1^H-NMR were deposited in the Biological Magnetic Resonance Databank under accession 27881. EPR data is available at doi: 10.17630/11b18167-18c0-42ae-a54c-1fe09432e91f. BED files of the TraDIS sequencing data are available at doi: 10.6084/m9.figshare.5280733. All other data has been published in the manuscript or as supporting information.

## ABBREVIATIONS

BEST: band-selective short transient excitation
DIPSI: decoupling in the presence of scalar interactions
DTT: dithiothreitol
EDTA: ethylene diamine tetra-acetic acid
EPR: electron paramagnetic resonance
HSQC: heteronuclear single quantum coherence
ICP: inductively coupled plasma
IPTG: isopropyl-β-thiogalactoside
ITC: isothermal titration calorimetry
MS: mass spectrometry
NMR: nuclear magnetic resonance
NTA: nitrilotriacetic acid
OES: optical emission spectrometry
SUMO: small ubiquitin-like modifier
TCEP: tris(2-carboxyethyl)phosphine
TOCSY: total correlated spectroscopy
TROSY: transverse relaxation optimised spectroscopy
UPF: unidentified protein function
UV: ultraviolet

## ACKNOWLEDGEMENTS

We thank J. Cole, J. Green, A. Peacock and O. Daubney for advice and assistance. We thank Drs C. Stark, S. Baker, M. Thompson, H. El Mkami and A. Shah for technical assistance and members of the Henderson, Lund and Grainger labs for insightful discussions. TCS was funded by the Biotechnology and Biological Sciences Research Council (BBSRC) Midlands Integrated Integrative Biosciences Training Partnership (MIBTP). MA was funded by Jouf University. DH and MJ were funded by BBSRC grant BB/L019434/1. TK was funded by BBSRC grant BB/P009840/1. EHAA was supported by the Ministerio de Ciencia, Tecnología e Innovación del govierno de Colombia and the British Council. JEL thanks the Royal Society for a University Research Fellowship and the Wellcome Trust for the Q-band EPR spectrometer (099149/Z/12/Z). NMR work was supported by the Wellcome Trust (099185/Z/12/Z), and we thank HWB-NMR at the University of Birmingham for providing open access to their Wellcome Trust-funded 900 MHz spectrometer.

