## Supporting Data for "Iron is a ligand of SecA-like metal-binding domains *in vivo*"

**Supporting figure S1.**

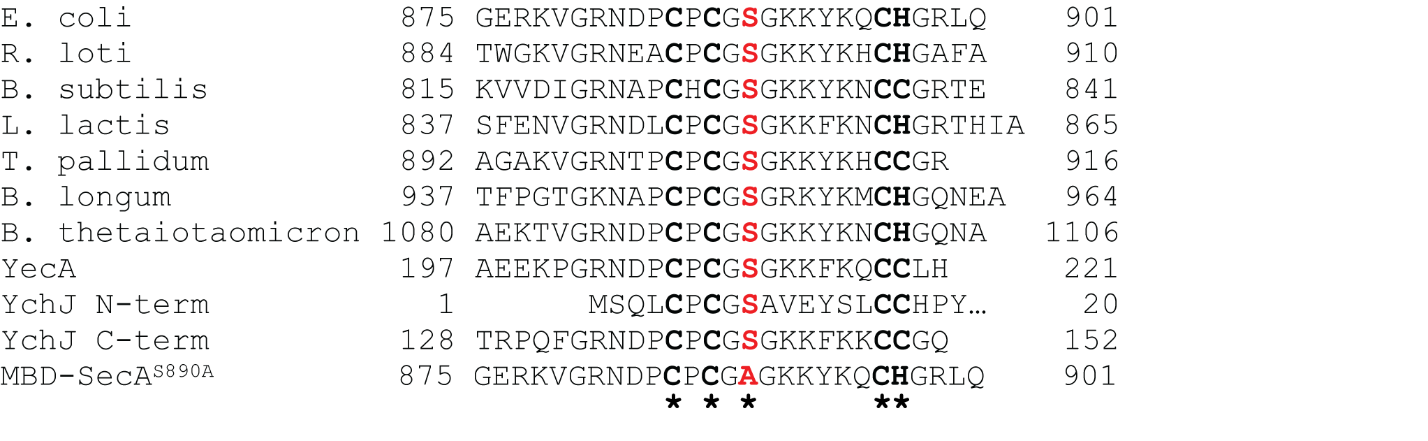

**Supporting figure S1. Sequence analysis of SecA-like MBDs.** Sequence alignment of the MBDs of SecA from *Escherichia coli*, *Rhizobium loti*, *Bacillus subtilis*, *Lactococcus lactis*, *Treponema pallidum*, *Bifidobacterium longum* and *Bacteriodes thetaiotamicron*, YecA from *Escherichia coli*, the N- and C-terminal MBDs of YchJ from *Escherichia coli* and the MBD-SecA^S889A^ peptide. The conserved metal-coordinating amino acids at positions 885, 887, 896 and 897 in *E. coli* SecA are bolded in black and starred. The conserved serine residue at position 889 in *E. coli* SecA is bolded in red.

**Supporting figure S2.**

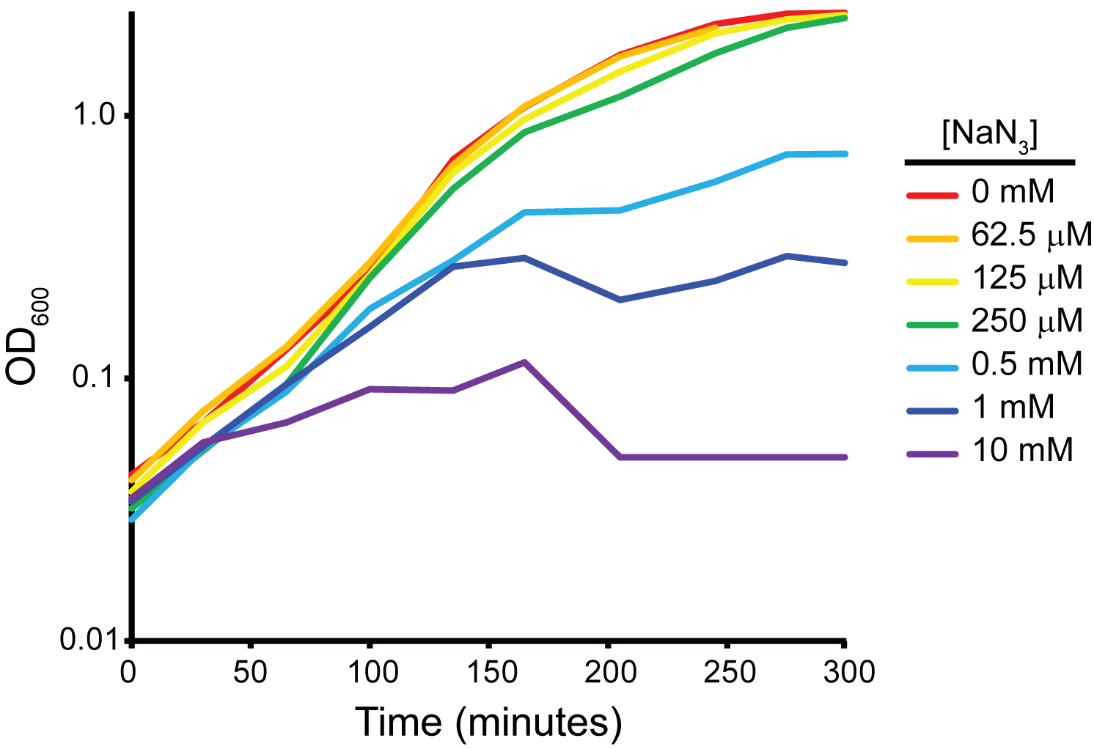

**Supporting figure S2. Effect of azide on the growth of *E. coli* BW25113.** *E. coli* strain BW25113 was grown in LB in the absence (red) or presence of 62.5 μM (orange), 125 μM (yellow), 250 μM (green), 500 μM (blue), 1 mM (indigo) or 10 mM (violet) sodium azide. Growth of the culture was monitored by optical density at 600 nm.

**Supporting figure S3.**

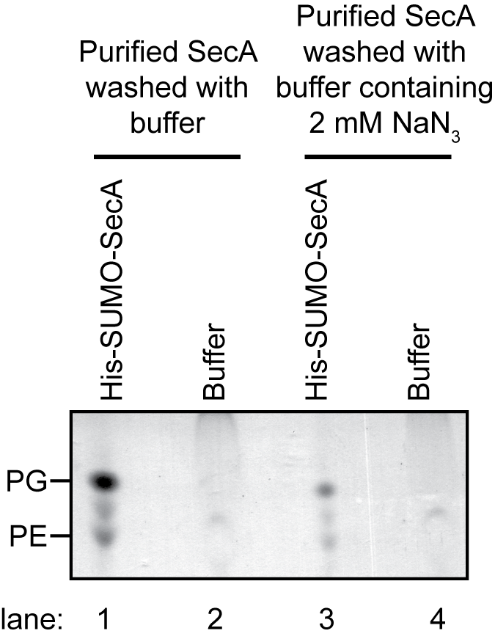

**Supporting figure S3. Effect of azide in wash buffers on the release of phospholipids by SecA.** His-SUMO-SecA was purified from untreated cells as in (A) and bound to a Ni-NTA beads. The beads were then washed with buffer containing (lanes 3 & 4) or lacking (lanes 1 & 2) 2 mM sodium azide. The lipids from 0.2 mg of protein were extracted from the eluted protein into 100 μl chloroform, and 20 μl of the extracted lipid (lanes 1 & 3) and the wash buffer (lanes 2 & 4) were resolved using TLC. The positions of PG, PE and a non-specific band from the wash buffer (*) are indicated.

**Supporting figure S4.**

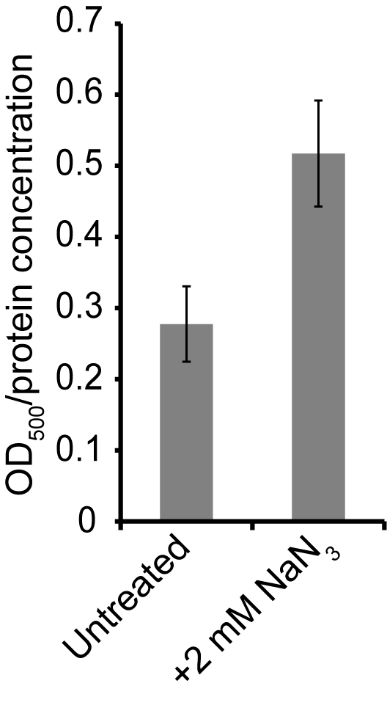

**Supporting figure S4. Effect of azide on the propensity of SUMO-CTT to aggregate.** Cells producing SUMO-CTT were incubated in the absence (untreated) or presence (+N_3_) of 2 mM NaN_3_ for 10 minutes. SUMO-CTT was purified from the cell lysates using streptactin beads and washed extensively with buffer. The bound protein was eluted from the streptactin beads using 7M guanidinium and was then diluted into buffer lacking guanidinium. Aggregation of the protein was measured using light scattering at 500 nm. Confidence intervals are the s.e.m.

**Supporting figure S5.**

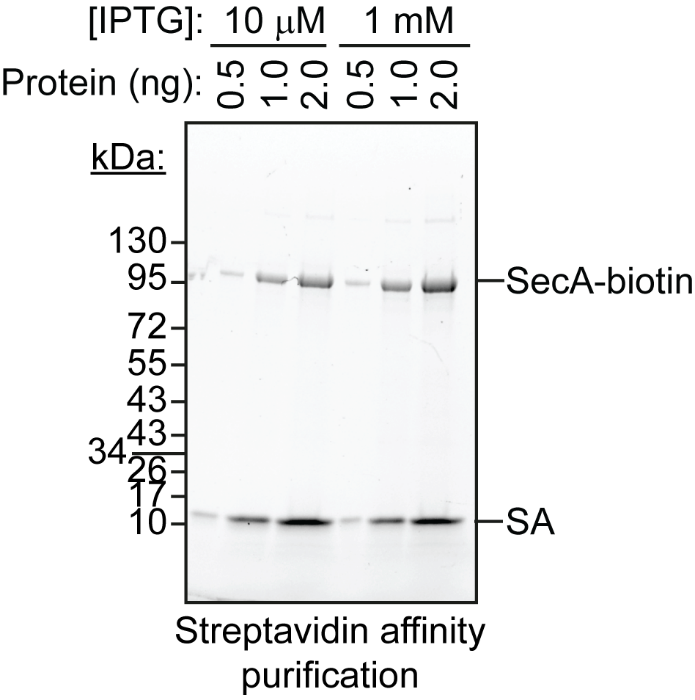

**Supporting figure S5. Purification of SecA-biotin from cell lysate using streptavidin.** DRH839 cells (Δ*secA* p*_trc_*-*secA-biotin*) were grown in the presence of 10 μM or 1 mM IPTG, as indicated, and SecA-biotin was purified from the cells using streptavidin-coated sepharose beads. After elution of the bound metal protein as described in figure 6, the protein was eluted from the beads by boiling in 1X Laemmli buffer, and the protein concentration in the samples was determined using Bradford. The indicated amount of protein was loaded on SDS-PAGE gel and visualised using Bio-Rad Stain-free technology. The molecular weights of a protein standard in kDa are indicated to the left. SA; streptavidin.

**Supporting figure S6.**

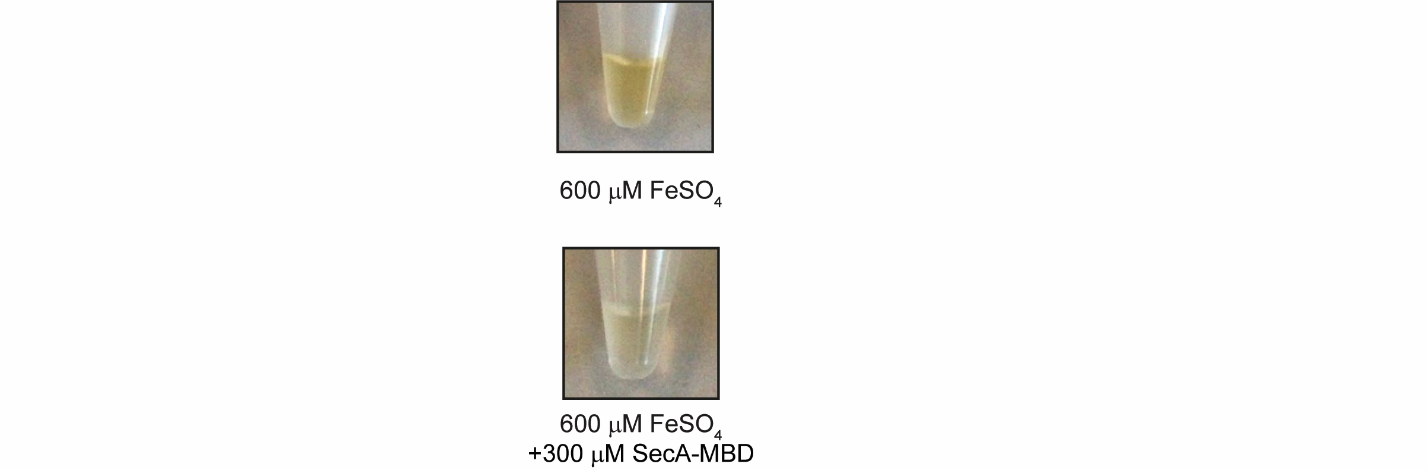

**Supporting figure S6. SecA-MBD peptide protects FeSO_4_ from formation of iron compounds.** 600 μM FeSO_4_ was incubated in the absence (above) or presence (below) of a 300 μM solution of a synthetic peptide consisting of the C-terminal 27 amino acids of SecA (SecA-MBD) and incubated at room temperature for 15 minutes.

**Supporting figure S7.**

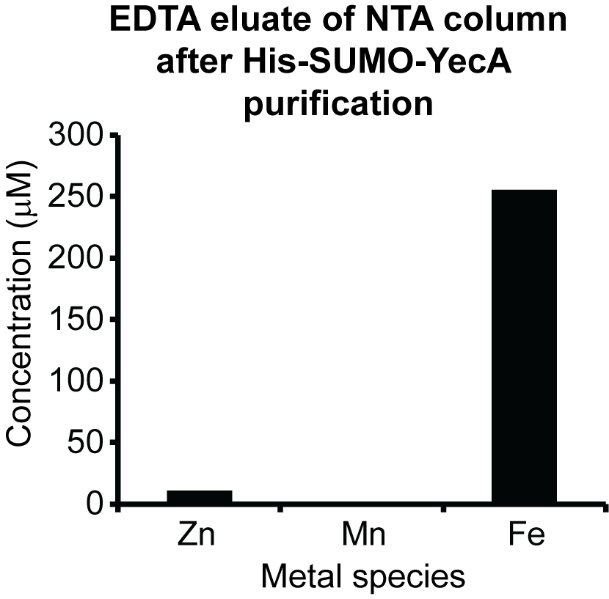

**Supporting figure S1. Analysis of the EDTA eluate from columns used to purify YecA.** Lysates of BL21(DE3) producing His-SUMO-YecA were passed over a Ni-NTA column and washed extensively with buffer containing 50 mM imidazole and 1 mM TCEP. Because the column retained a yellow colour after elution of the bound protein, the residual metal was stripped from the Ni-NTA column using 1 mM EDTA. The zinc, manganese and iron content of the EDTA eluate was then determined using ICP-OES.

**Supporting figure S8.**

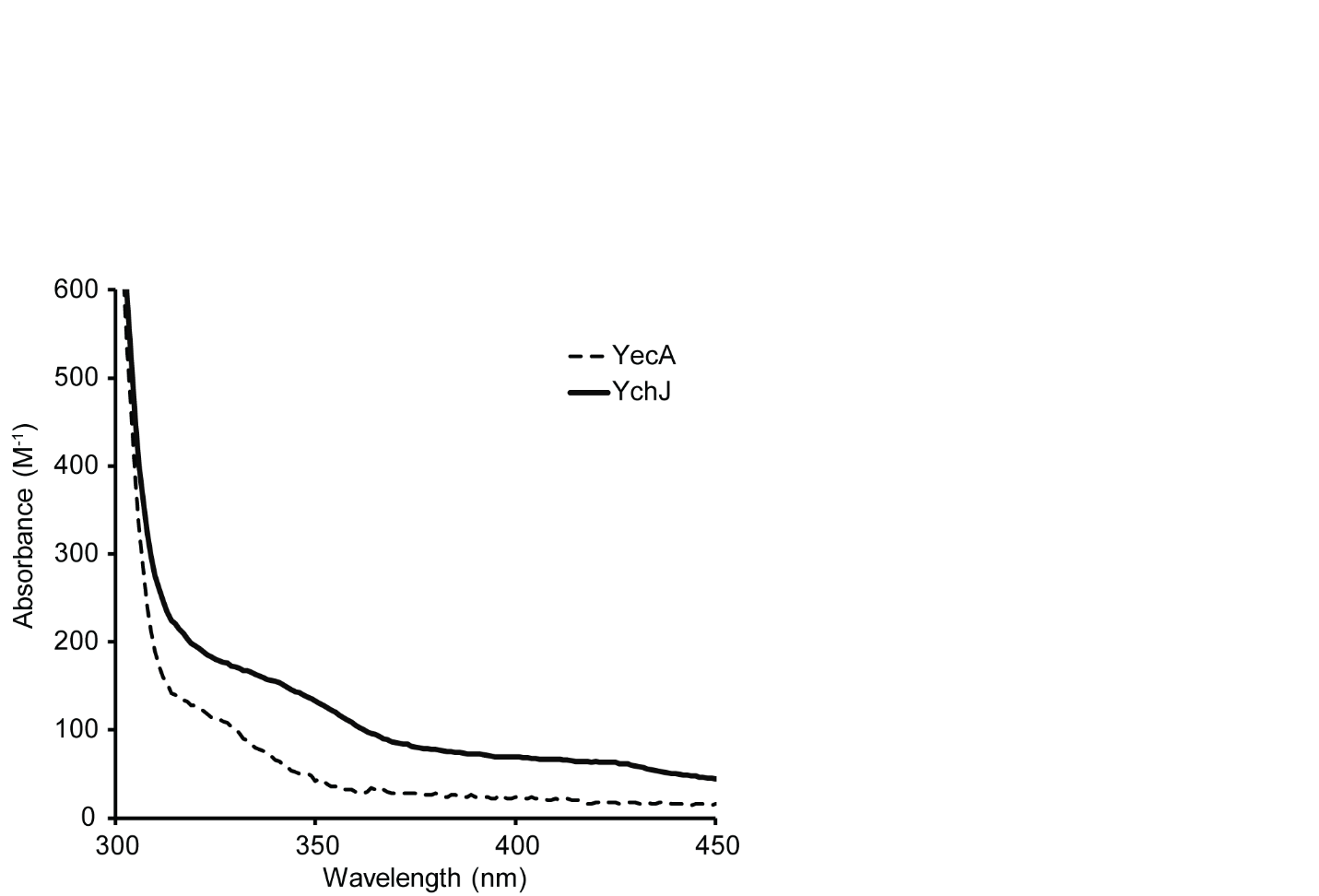

**Supporting figure S8. Absorbance spectra of purified YecA and YchJ.** Buffer subtracted absorbance spectra YecA (dashed) and YchJ (solid black). The absorbance at each wavelength was normalised to the concentration of the protein and the number of MBDs in each protein (*i.e.* the molar absorbance spectrum of YchJ was divided by two).

**Supporting figure S9.**

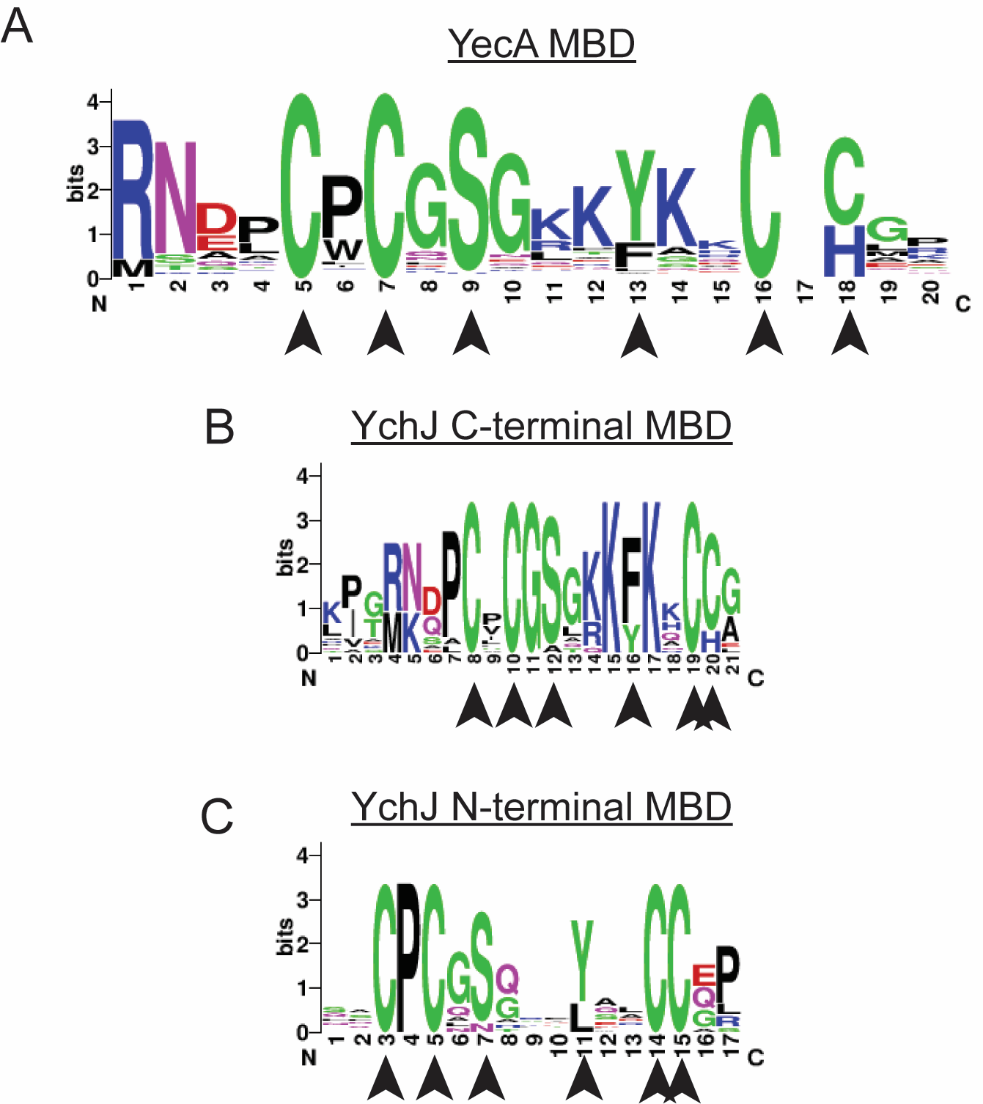

**Supporting figure S9. Consensus sequences of the YecA and YchJ MBDs.** Logos representing the consensus sequences of the YecA MBD (A), the C-terminal YchJ MBD (B) and the N-terminal YchJ MBD (C) were constructed from the alignments of 10 sequences for YecA, 16 sequences for the YchJ C-terminal MBD and 15 sequences for the YchJ N-terminal MBD from NCBI.

**Supporting figure S10.**

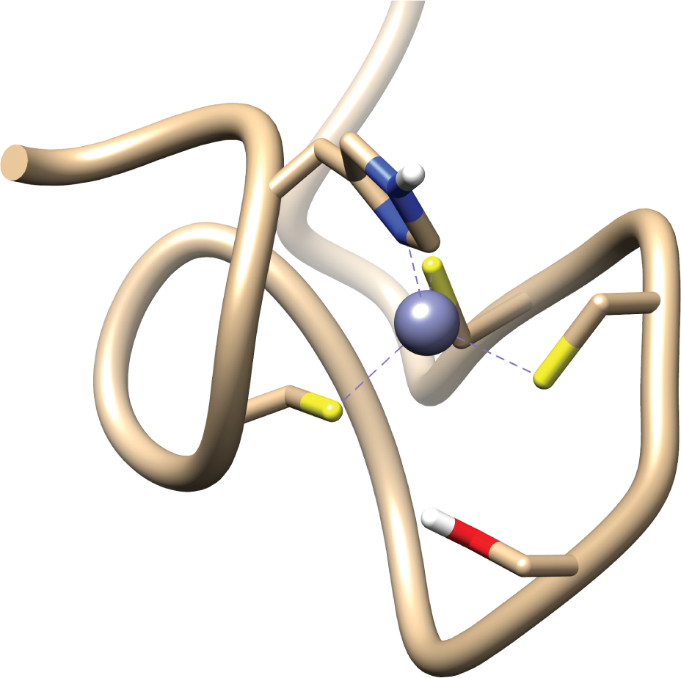

**Supporting figure S10. Structure of the SecA MBD bound to zinc.** Ribbon diagram of an example structural models of the SecA MBD from PDB file 1SX1 (7). The side-chains of the metal-coordinating amino acids and conserved Ser-889 are depicted as sticks (yellow, sulfur; blue, nitrogen; red, oxygen). Image was rendered using UCSF Chimera v. 1.12 (48).

**Supporting figure S11.**

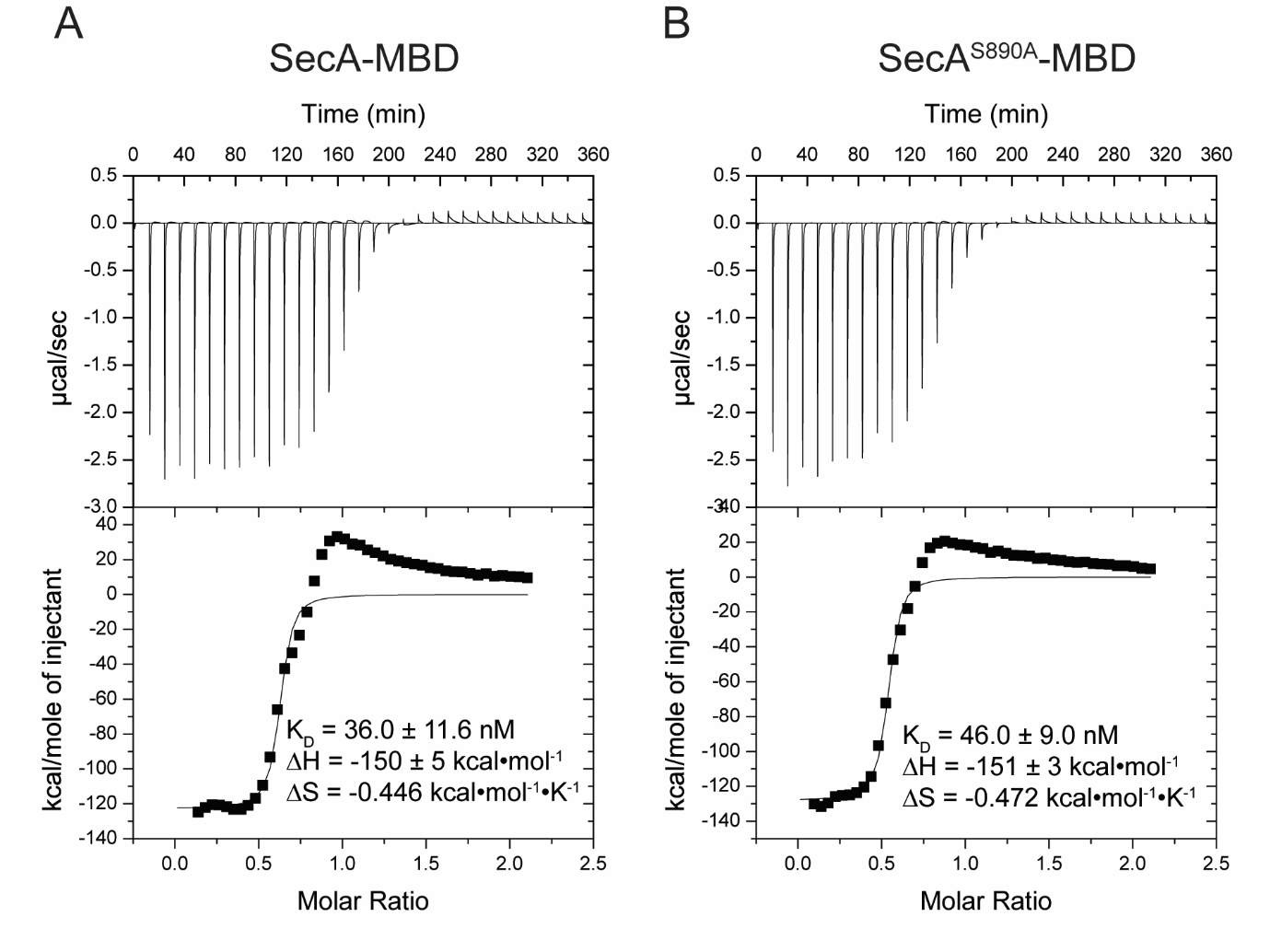

**Supporting figure S11. Determination of the affinity of SecA-MBD and SecA-MBD^S889A^ for Zn^2+^ by ITC.** The heat exchange upon mixing solutions containing SecA-MBD peptide (A) and SecA-MBD^S889A^ peptide (B) with ZnSO_4_ was measured by isothermal titration calorimetry (ITC; above). K_D_, ΔH and ΔS determined from fits of the heat exchange curves (below).

**Supporting figure S12.**

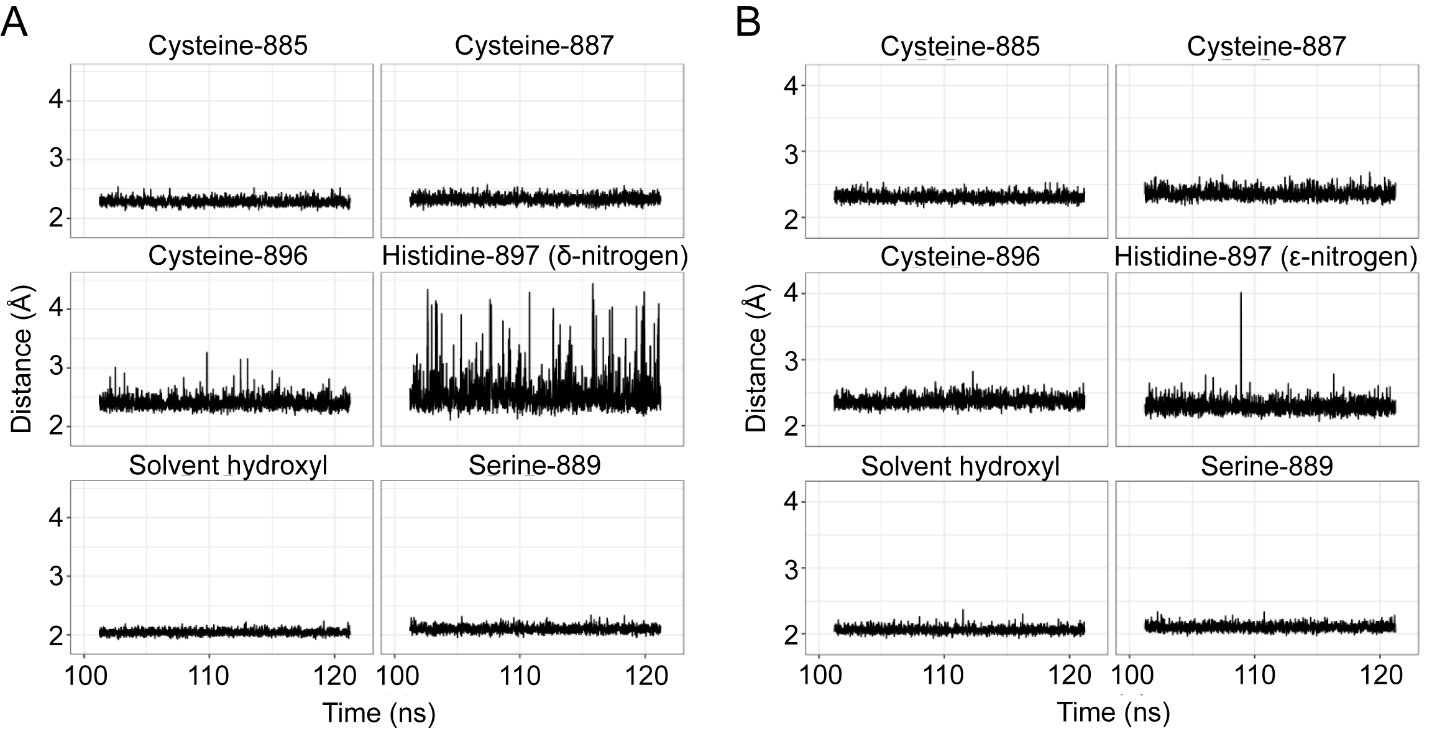

**Supporting figure S12. Distance measurements of metal-coordinating atoms from molecular dynamics simulations of SecA-MBD octahedrally bound to Fe^2+^.** Depicted are the distances between the metal ion and the metal-coordinating atoms in Cys-885, Cys-887, Ser-889, Cys-896, His-897 and the solvent hydroxyl ion recorded at 10 ps intervals during the final 20 ns of a 100 ns simulation. (A) Continuation with reduced restraints of the 100 ns simulation with His-897 restrained *via* the δ-nitrogen. (B) Continuation with reduced restraints of the 100 ns simulation with His-897 restrained *via* the ε-nitrogen.

**Supporting table S1. Effect of transition metal availability on azide sensitivity**

| **Condition** | **Zone of clearing*** |
| --- | --- |
| LB | 27.3 ± 2.5 mm |
| LB + 1 mM EDTA | 33.3 ± 2.4 mm |
| LB + 100 μM ZnSO_4_ | 28.5 ± 1.5 mm |
| LB + 100 μM FeSO_4_ | 24.5 ± 0.5 mm |

*Diameter of the zone clearing in a lawn of *Escherichia coli* MC4100 surrounding a 6 mm antibiotic assay filter disc containing 10 μl of 1 M sodium azide.

**Supporting table S2. Atomic charges used in molecular dynamics simulations for deprotonated cysteine (CYM) and deprotonated serine (SEO).**

| **CYM** | | **SEO** | |
| --- | --- | --- | --- |
| **Atom** | **Charge** | **Atom** | **Charge** |
| N | -0.415700 | N | -0.415700 |
| H | 0.271900 | H | 0.271900 |
| CA | -0.035100 | CA | -0.035100 |
| HA | 0.050800 | HA | 0.050800 |
| CB | -0.241300 | CB | -0.191300 |
| HB2 | 0.112200 | HB2 | 0.112200 |
| HB3 | 0.112200 | HB3 | 0.112200 |
| SG | -0.884400 | OG | -0.934400 |
| C | 0.597300 | C | 0.597300 |
| O | -0.567900 | O | -0.567900 |

**Supporting table S3**. **Summary of different restraint parameters applied during simulations.**

| **Restraint** | **r_1_^†^** | **r_2_^†^** | **rk_2_^†^** | **r_3_^†^** | **r_4_^†^** | **rk_3_^†^** |
| --- | --- | --- | --- | --- | --- | --- |
| Between the metal ion and the coordinating atoms | 1.30 Å | 1.80 Å | 0 kcal/(mol•Å) | 2.30 Å | 2.80 Å | 10 kcal/(mol•Å) |
| Between adjacent atoms on vertices of the octahedron (see table S3) | 2.05 Å | 2.55 Å | 0 kcal/(mol•Å) | 3.25 Å | 3.75 Å | 2.5 kcal/(mol•Å) |
| Torsional restraint^*^ on Cys-885:S, Cys-887:S, Cys-896:S and His-897:N(D/E) | -7˚ | -5˚ | 10 kcal/(mol•rad) | 5˚ | 7˚ | 10 kcal/(mol•rad) |

**^†^**The restraints define an energy well that has a flat bottom when the restrained distance is between r_2_ and r_3_, i.e. no restraint force is applied within bounds, a parabolic shape between r_2_ and r_1_, and between r_3_ and r_4_, and is linear when lower than r_1_ or greater than r_4_, with the force constants rk_2_ and rk_3_, respectively. For more information, the Amber 18 manual can be consulted (Case, D. A., Ben-Shalom, I. Y., Brozell, S. R., Cerutti, D. S., Cheatham III, T. E., Cruzeiro, V. W. D., ... & Gohlke, H. AMBER 18. 2018. *San Francisco: University of California.*)

^*^Even though the force constant for an angle restraint has units kcal/(mol•rad), the values of r_x_ are still given in degrees of arc.

**Supporting table S4. List of restraints between adjacent atoms on vertices of the octahedron.**

| His-897:N(D/E) | Cys-887:S |
| --- | --- |
| His-897:N(D/E) | Cys-896:S |
| His-897:N(D/E) | O of hydroxyl |
| His-897:N(D/E) | Ser-889:OG |
| Cys-887:S | Cys-885:S |
| Cys-887:S | O of hydroxyl |
| Cys-887:S | Ser-889:OG |
| Cys-885:S | Cys-896:S |
| Cys-885:S | O of hydroxyl |
| Cys-885:S | Ser-889:OG |
| Cys-896:S | O of hydroxyl |
| Cys-896:S | Ser-889:OG |

List of pairs of atoms that are adjacent on the surface of the octahedron. Each pair was subjected to a distance restrain of 2.5 kcal/(mol•Å), as defined in table S2.

**Supporting data S1. Analysis of transposon insertion mutants in TraDIS library after outgrowth in LB or LB containing NaN_3_.** Excel spreadsheet detailing: (A) Sheet 1. Number of insertions in each gene in the *E. coli* genome after growth of the TraDIS library in LB, LB containing 0.25 mM NaN_3_ or LB containing 0.5 mM NaN_3_. Genomic DNA from one experiment was prepared and sequenced twice to increase the number of sequencing reads and to examine the reproducibility of the results. For each run the number of insertions in each gene was normalised to the total number of sequencing reads. (B) Sheet 2. The degree of enrichment or depletion of insertion mutations in the library after growth in the presence of azide was calculated by determining the log (base 2) of the ratio of the number of insertions in a gene after outgrowth in the azide-treated conditions to the number of insertions after outgrowth in LB. The BED files used to derive the data presented in parts A and B is available at figshare.com (doi: 10.6084/m9.figshare.5280733).

**Supporting data S2. *E. coli* proteins containing CXCX_N_CH or CXCX_N_CH motifs.** Table containing UniProt accession number, UniProt protein name, the CXCX_N_CH- or CXCX_N_CH-containing amino acid sequence and the identity of any coordinated metals (if known) of all proteins in *E. coli* that contain the sequence CXCX_N_CH- or CXCX_N_CH.

**Supporting data S3. Structure file of SecA MBD coordinating a hexavalent metal *via* the δ-nitrogen of His-897.** File contains 10 structures from the last 1 ns of a 100 ns simulation.

**Supporting data S4. Structure file of SecA MBD coordinating a hexavalent metal *via* the ε-nitrogen of His-897.** File contains 10 structures from the last 1 ns of a 100 ns simulation.
